# SEC is an anti-angiogenic virulence factor that promotes *Staphylococcus aureus* Infective Endocarditis Independent of Superantigen Activity

**DOI:** 10.1101/2019.12.13.875633

**Authors:** Kyle J. Kinney, Sharon S. Tang, Xiao-Jun Wu, Phuong M. Tran, Nikhila S. Bharadwaj, Katherine N. Gibson-Corley, Ana N. Forsythe, Katarina Kulhankova, Jenny E. Gumperz, Wilmara Salgado-Pabón

## Abstract

The superantigen staphylococcal enterotoxin C (SEC) is critical for *Staphylococcus aureus* infective endocarditis (SAIE) in rabbits. Superantigenicity, its hallmark function, was proposed to be a major underlying mechanism driving SAIE but was not directly tested. With the use of *S. aureus* MW2 expressing SEC toxoids, we show that superantigenicity does not sufficiently account for vegetation growth, myocardial inflammation, and acute kidney injury in the rabbit model of native valve SAIE. These results highlight the critical contribution of an alternative function of superantigens to SAIE. In support of this, we provide evidence that SEC exerts anti-angiogenic effects by inhibiting branching microvessel formation in an *ex vivo* rabbit aortic ring model and by inhibiting endothelial cell expression of one of the most potent mediators of angiogenesis, VEGF-A. SEC’s ability to interfere with tissue re-vascularization and remodeling after injury serves as a mechanism to promote SAIE and its life-threatening systemic pathologies.

## Introduction

*Staphylococcus aureus* is the leading cause of infective endocarditis (IE) in high-income countries (*1–3*). *S. aureus* IE (SAIE) is an acute and invasive infection of the cardiac endothelium characterized by the appearance of vegetative lesions that form predominantly on heart valves (*4*). The pathognomonic vegetations are a meshwork of bacterial aggregates and host factors such as fibrin, fibrinogen, platelets, and erythrocytes that form predominantly on heart valves (*5*). SAIE results in significant damage to cardiac structures, in particular the valves and myocardium, due to tissue toxicity and necropurulent inflammation (*6*). Once established, SAIE can lead to severe complications, most notably congestive heart failure, stroke, acute kidney injury, and septic shock (*3, 6, 7*). Treatment of SAIE is challenging, requiring prolonged antibiotic therapy or surgery to remove infected valves (*3, 6*). Even with treatment, SAIE has a high rate of recurrence and a 22-66% mortality rate (*1, 3*). Infections are frequently associated with methicillin-resistant *S. aureus* (MRSA) which complicate treatment and increase risk of mortality (*8*). Furthermore, life-saving medical interventions (e.g., valve replacement, cardiac devices, and hemodialysis), an increasing population with underlying conditions (e.g., diabetes mellitus and immunosuppression), and advanced age also increase the risk of acquiring SAIE (*1, 3*). As a result, the incidence of SAIE has continued to increase (*3*). Unfortunately, the advances in cardiovascular medicine achieved in the last decade have failed to improve SAIE outcomes. Thus, understanding the mechanisms driving SAIE pathophysiology is not only of fundamental interest, particularly as it relates to bacterial factors critical for vegetation formation and secondary complications, but also of utmost importance to develop effective intervention strategies.

Epidemiological studies demonstrate a strong association between SAIE and a select group of superantigen (SAg) genes, where 18-25% of SAIE clinical isolates encode *entC* (staphylococcal enterotoxin C; SEC), 9-20% encode *tstH* (toxic shock syndrome toxin; TSST1), and 58-90% encode the enterotoxin gene cluster (*egc*) (*9*). Consistent with these studies, SEC, TSST1, and the *egc* SAgs SEI, SEM, SEO, and SE *like* (*l*)-U all contribute to IE and metastatic infection in experimental IE (*10, 11*). However, the underlying mechanism by which SAgs contribute to SAIE pathogenesis remains speculative. Classically, SAgs are known for their potent T cell mitogenic activity resulting in dysregulated activation and cytokine production leading to inflammatory syndromes, and in extreme cases, toxic shock (*12*). Superantigenicity results from toxin cross-linking of the Vβ-chain of the T-cell receptor (TCR) to the major histocompatibility complex class II (MHC-II) receptor on antigen presenting cells (*12*). Of relevance to SAIE, endothelial cells also express MHC-II and function as conditional antigen presenting cells capable of cross-linking Vβ–TCR resulting in endothelium-mediated superantigenicity (*13*).

The dysregulated immune activation caused by SAgs distracts and diverts the immune system (*14*). It is also the etiology of multiple diseases including atopic dermatitis, pneumonia, extreme pyrexia, purpura fulminans, and toxic shock syndrome (*12*). The commonly accepted model of the role of SAgs in SAIE includes localized or systemic superantigenicity that causes dysregulation of the immune system preventing clearance of *S. aureus* from the infected heart endothelium. SAgs also cause capillary leak and hypotension which changes the hemodynamics of the vascular system (*12*). This alteration of blood flow may enhance vegetation formation. However, the requirement of superantigenicity in the pathogenesis and pathophysiology of SAIE has not been directly tested. In this study, we addressed the hypothesis that superantigenicity promotes SAIE and disease sequelae.

We used the rabbit model of native valve IE with the well-characterized methicillin-resistant *S. aureus* strain MW2 (SEC^+^) and MW2 stably expressing SEC toxoids (TCR or MHC-II/TCR inactivated) to provide evidence that SEC critically contributes to vegetation growth, the magnitude of myocardial inflammation, and injury to the renal and hepatic systems independent of superantigenicity. We demonstrate that development of septic vegetations are a pre-requisite for decreased renal function, while superantigenicity resulting from SAIE exacerbates hepatocellular injury and systemic toxicity.

With the use of the *ex vivo* rabbit aortic ring model of angiogenesis and human aortic endothelial cells, we provide evidence that SEC inhibits re-vascularization and inhibits expression of Vascular Endothelial Growth Factor (VEGF)-A, a potent pro-angiogenic factor. These results demonstrate that SEC is an antiangiogenic virulence factor that directly modifies endothelial cell function by a mechanism that can promote SAIE and its associated systemic pathologies.

### *S. aureus* MW2 expressing SEC toxoids are deficient in superantigen activity *in vitro* and *in vivo*

SAg activity is the most potent biological function of staphylococcal enterotoxins described to date and causes lethal pathologies. To establish whether superantigenicity promotes development of SEC-mediated SAIE, we complemented *S. aureus* MW2Δ*sec* to produce SEC with an inactive T-cell receptor (TCR)-binding site (SEC_N23A_) or to produce SEC with dual inactivation of the TCR- and MHC class II-binding sites (SEC_N23A/F44A/L45A_) (Table S1-S2). Due to its biological inactivation, SEC_N23A_ and mutations thereof are excluded from the CDC select agent toxins (*15*). Importantly, several vaccination studies have shown no disruption in SEC_N23A_ toxin structure and in the antigenic nature of the protein (*16–19*). MHC-II is expressed by non-hematopoietic cells such as epithelial cells and endothelial cells (*13*). Hence, to exclude the possibility that interactions with MHC-II may promote disease, we introduced two mutations (F44A/L45A) into SEC_N23A_ previously shown to effectively inactivate MHC-II binding (*18–21*). *S. aureus* expressing SEC_N23A_ or SEC_N23A/F44A/L45A_ were confirmed to produce SEC toxoids at similar levels as wildtype SEC (Fig. S1A and B) and to exhibit no defects in growth in liquid culture (Fig. S1C) or defects in hemolytic activity as proxy for assessing functionality of the Agr system (Fig. S1D and E).

To provide evidence that *S. aureus* expressing toxoids are deficient in SEC superantigenicity, cell-free supernates from overnight cultures of MW2 (*S. aureus* SEC^+^), the isogenic deletion strain MW2Δ*sec* (*S. aureus* SEC^KO^), and *S. aureus* expressing toxoids (*S. aureus* SEC_N23A_ and *S. aureus* SEC_N23A/F44A/L45A_) were tested in a T cell proliferation assay. For this, human peripheral blood mononuclear cells (PBMCs) from 3 donors were cultured in the presence of 0.3, 2, or 10-μl volumes of supernates and the replication index of CD4^+^ T cells calculated 6 days post-exposure. Purified SEC (10 μg) was used as proliferation control. As expected, *S. aureus* SEC_N23A_ and *S. aureus* SEC_N23A/F44A/L45A_ were both deficient in SEC superantigenicity as T cell proliferation was reduced to levels at or below those resulting from *S. aureus* SEC^KO^ (Fig. 1A and Fig. S2). Similarly, the serum from rabbits infected with *S. aureus* SEC_N23A_ or SEC_N23A/F44A/L45A_ in the native valve IE model exhibited significantly decreased concentrations of IL-6 and lactate dehydrogenase (LDH) compared to those infected with *S. aureus* SEC^+^ (Fig. 1B and C). IL-6 is a cytokine produced at high levels as a result of superantigen activity (*22*). LDH, found in almost every cell in the body, is released into the systemic circulation as a result of tissue injury including the cytotoxic effects of cytokine storms (*23–25*). On average, both IL-6 and LDH levels in rabbits infected with *S. aureus* expressing toxoids were not significantly different than those in rabbits infected with *S. aureus* SEC^KO^ (Fig. 1B and C). These results are consistent with inactivation of superantigen activity in SEC_N23A_ and SEC_N23A/F44A/L45A_ and provide further evidence of deficient superantigen activity of *S. aureus* expressing toxoids.

**Figure 1.**
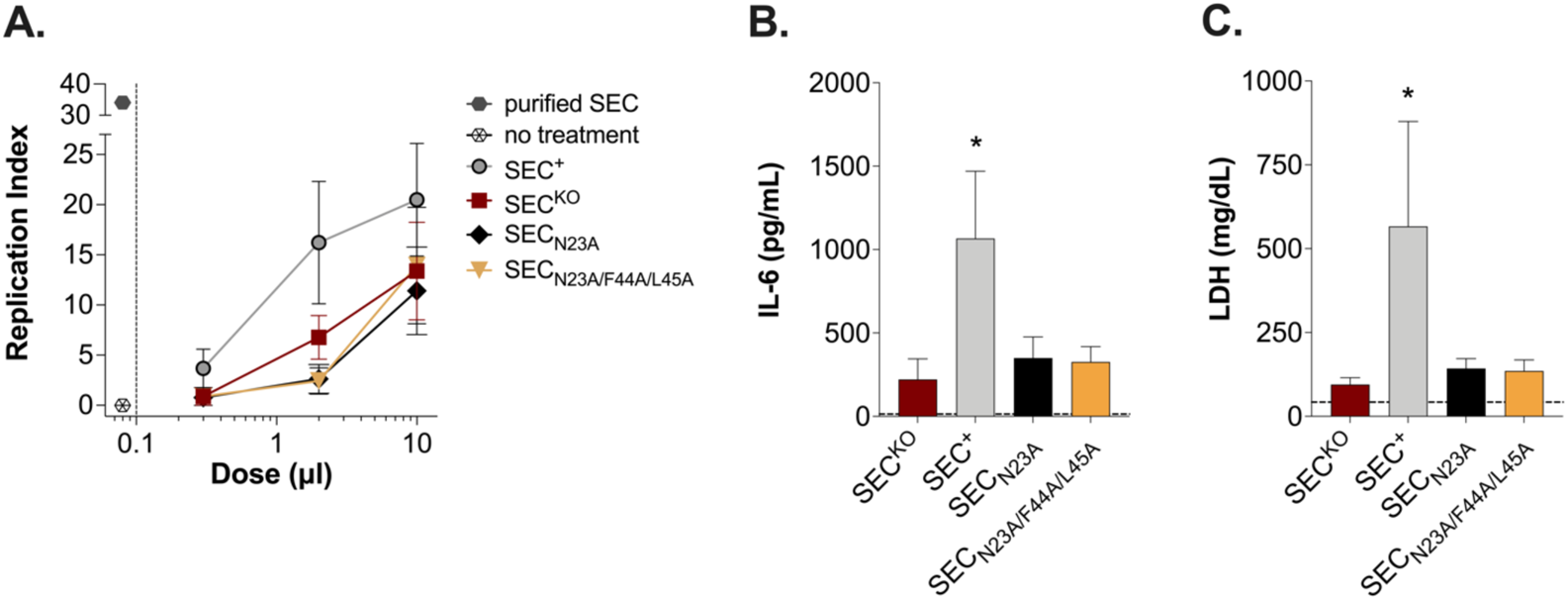
SEC toxoids are deficient in superantigen activity. (**A**) CD4^+^ T cell replication index of peripheral blood mononuclear cells (PBMCs) stimulated for 6 days with 0.3, 2, and 10 μl of cell-free supernates from overnight cultures of *S. aureus* SEC^KO^, SEC^+^, SEC_N23A_, and SEC_N23A/F44A/L45A_. PBMCs ± 10 μg of purified SEC were used as controls. (**B-C**) Serum levels of interleukin-6 (IL-6) and lactate dehydrogenase (LDH) 48-96 h post-infection with indicated *S. aureus* strains in experimental infective endocarditis. *S. aureus* SEC^+^ (n=12), *S. aureus* SEC^KO^ (n=15), *S. aureus* SEC_N23A_ (n=17), and *S. aureus* SEC_N23A/F44A/L45A_ (n=17). Data are represented as mean (± SEM). The dashed line is the average analyte value of all rabbits pre-infection. Statistical significance was determined by one-way ANOVA with Holm-Šídák’s multiple comparisons test with each SEC-producing strain compared to *S. aureus* SEC^KO^. All experimental groups were tested against pre-infection values and were statistically significant, *p* < 0.0001. (**B**) *, *p* = 0.03. (**C**) *, *p* = 0.02.

### Superantigenicity is not sufficient to promote vegetation formation and high lethality

*S. aureus* SEC^+^, *S. aureus* SEC^KO^, *S. aureus* SEC_N23A_, and *S. aureus* SEC_N23A/F44A/L45A_ were tested in the rabbit native-valve IE model. Strains were inoculated intravenously with 2 – 4 x10^7^ total CFU after mechanical damage to the aortic valve and monitored for a period of 4 days. During that period, rabbits infected with *S. aureus* SEC^KO^ had a ~66% decrease in overall vegetation formation where 8/12 rabbits had no vegetations (Fig. 2A and B) and 4/12 had very small vegetations (sent to pathology) with an estimated weight of 5 – 15 mg. In stark contrast, *S. aureus* SEC^+^ formed vegetations in nearly all rabbits (14/15). Vegetation size ranged from 11 – 116 mg with most vegetations weighing >25 mg (Fig. 2A and B). Six vegetations were sent for histopathology by a board-certified veterinary pathologist. Surprisingly, complementation with SEC_N23A_ or SEC_N23A/F44A/L45A_ restored vegetation formation to wild type levels in most rabbits, with vegetation sizes ranging from 12 – 103 mg for SEC_N23A_ (in 13/17 rabbits) and 3 – 107 mg for SEC_N23A/F44A/L45A_ in 12/17 rabbits (Fig. 2A and B).

**Figure 2.**
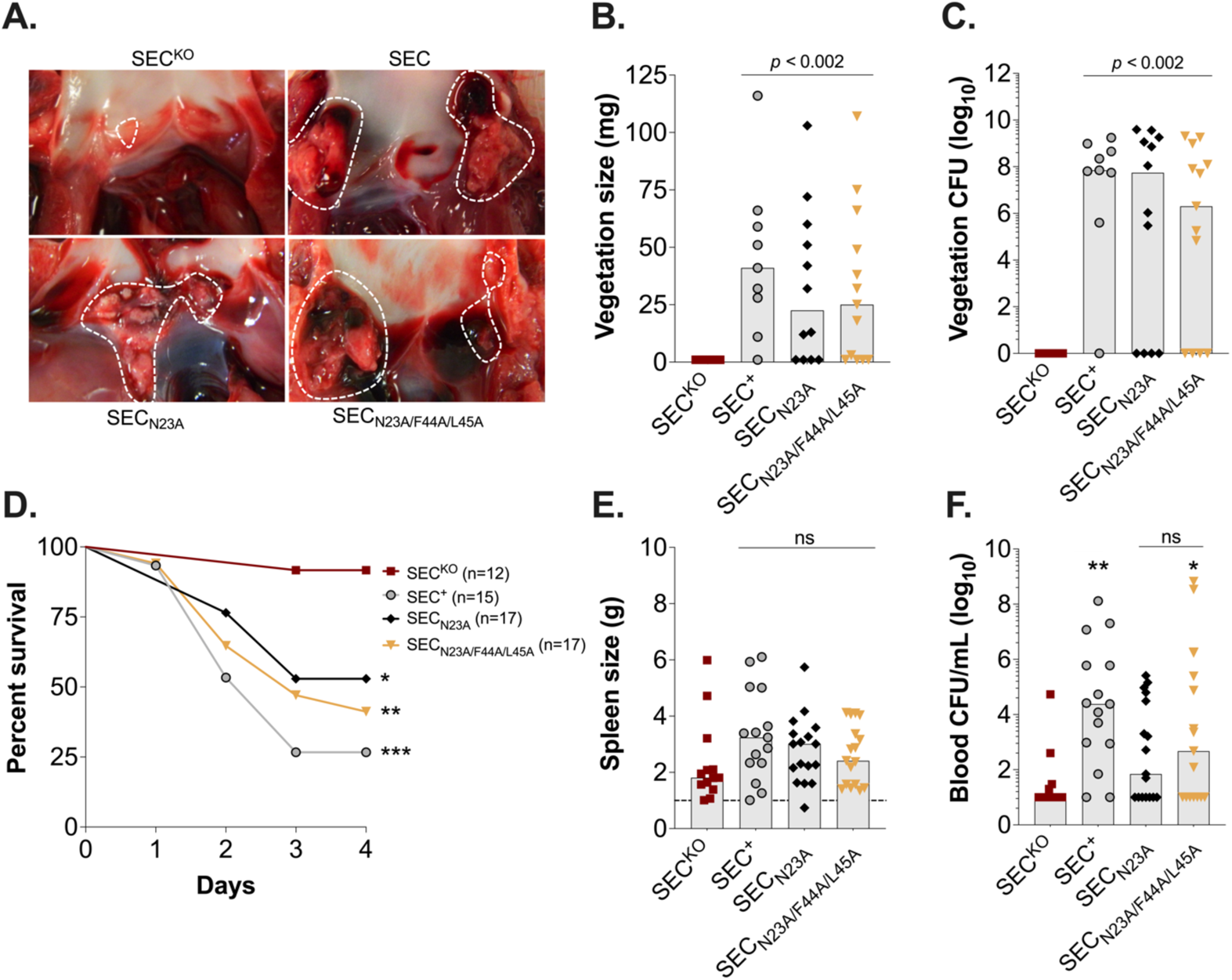
SEC is required for vegetation formation independent of superantigenic activity in the rabbit model of native valve infective endocarditis and sepsis. (**A**). Representative images of aortic vegetations (areas indicated with dash-lines). (**B**) Total weights of vegetations dissected from aortic valves from infected rabbits. (**C**) Bacterial counts recovered from aortic vegetations. (A-C) Bars represent median value. Statistical significance, one-way ANOVA with Holm-Šídák’s multiple comparisons test compared to SEC^KO^. (**D**) Percent survival of rabbits infected intravenously with 2×10^7^–4×10^7^ CFU of indicated strain over 4 days. *, *p* = 0.03, **, *p* = 0.006, ***, *p* = 0.0007 log-rank Mantel-Cox Test. (**E**) Enlargement of the spleen resulting from *S. aureus* infection. (Dashed line) Average spleen size of uninfected rabbit controls (0.86 ± 0.1, n = 3). (**F**) Bacterial counts per milliliter of blood recovered from rabbits *post-mortem*. (**D-F**) Bars represent median value. *, *p* = 0.03, **, *p* = 0.003, ns = not significant, one-way ANOVA with Holm-Šídák’s multiple comparisons test. Statistical significance compared to SEC^KO^.

Altogether, infection with *S. aureus* expressing toxoids produced vegetations in 74% of rabbits (25/34), compared to 93% (14/15) in rabbits infected with *S. aureus* SEC^+^. Vegetations formed by *S. aureus* SEC_N23A_ and *S. aureus* SEC_N23A/F44A/L45A_ also had bacterial counts comparable to the wildtype strain (Fig. 2C). Consistent with the contribution of vegetation formation to lethal systemic complications, infection with either *S. aureus* SEC_N23A_ or *S. aureus* SEC_N23A/F44A/L45A_ led to high lethality, with ~50% of rabbits succumbing to infection during the experimental period compared to 73% of rabbits infected with *S. aureus* SEC^+^ (Fig. 2D). Rabbits infected with *S. aureus* expressing toxoids also exhibited higher degrees of splenomegaly (Fig. 2E) and on average higher bacteremia (>1×10^3^ CFU/ml) than those infected with *S. aureus* SEC^KO^, although only those infected with SEC_N23A/F44A/L45A_ exhibited significantly higher bacteremia (Fig. 2F). Overall, there is an ~20% decrease in vegetation formation and lethality in rabbits infected with strains producing toxoids. Thus, superantigenicity alone does not fully account for the high lethal outcomes associated with SEC production in experimental SAIE. These results highlight a critical requirement of SEC in SAIE that is independent of superantigenicity and MHC class II interactions.

### Superantigenicity-independent effects drive myocardial inflammation in SAIE

SAIE presents as rapidly-growing vegetative lesions resulting in the quick destruction of valvular leaflets and progression of the infection into the myocardium and adjacent structures (*5*). We then asked whether SEC superantigenicity promotes extension of the vegetative lesion into the surrounding tissue changing the overall cardiac pathology. To address this, we performed histopathological analyses on transverse sections of hearts containing vegetations (Fig. S3 and S4, Table S3). Hearts from rabbits infected with *S. aureus* SEC^+^, *S. aureus* SEC_N23A_, or *S. aureus* SEC_N23A/F44A/L45A_ were selected randomly based on presence of vegetations (mean size 6.6 ± 2 mm^2^, n=15). We processed all hearts of *S. aureus* SEC^KO^ infected rabbits with visible vegetations (mean size 2.7 ± 1 mm^2^, n=4) and 3 hearts without vegetations. We recognize there is selection bias in this case (and the data will reflect that), but we wanted to learn pathologies associated with vegetation formation, including the 33% that forms in the absence of SEC production. Consistent with histopathology of SAIE described in humans, *S. aureus* vegetations in rabbits were composed of large aggregates of bacterial colonies interspersed in a fibrinous meshwork of host factors and cell debris (Fig. S3-S4). However, the vegetative lesions were heterogeneous across infection groups in presentation (location and size of bacterial clusters) and in the magnitude of suppurative intracardial complications (myocardial inflammation and septic coronary arterial emboli) (Fig. 3). In rabbits infected with *S. aureus* SEC^+^, most vegetations were located on aortic valve cusps and valve leaflets, with large clusters of bacteria present on the leaflets and intermixed within the central core of the vegetation adjacent to the aorta. A few vegetations formed on the aortic wall, were transmural (across the entire wall), and extended into the adjacent adipose tissue (Fig S4A). Rabbits infected with *S. aureus-*producing SEC toxoids (SEC_N23A_ or SEC_N23A/F44A/L45A_) exhibited very similar histologic presentation to each other and to those infected with *S. aureus* SEC^+^ (Fig S4B and C).

**Figure 3.**
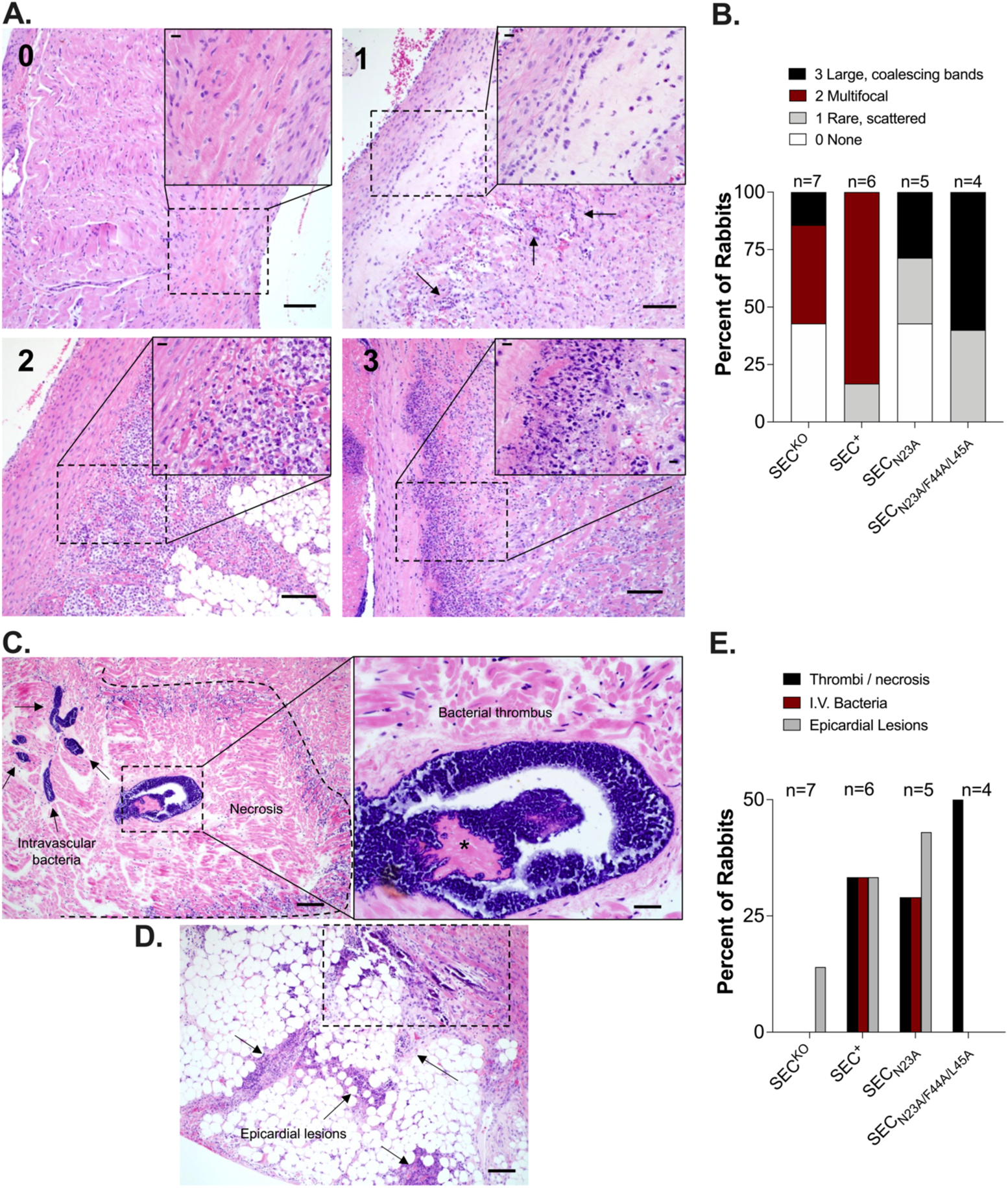
Vegetative lesions present with bacterial infiltration, inflammation, and necrosis. (**A**) Examples of histopathologic assessment of myocardial inflammation during rabbit model of native valves infective endocarditis (Graded 0-3). Inflammation was graded based on the amount of inflammatory cell infiltrate noted within the myocardium. 0 = no inflammation, 1 = rare, scattered infiltrate (arrows), 2 = multifocal bands of infiltrate, 3 = large, coalescing bands of diffuse infiltrate with necrosis. Bar = 100 µm, inset bar = 20 µm. (**B**) Scoring of myocardial inflammation from H&E-stained images 48 – 96 h post-infection. No statistical significance among wildtype and toxoid groups, Fisher’s exact test with categorical pathology grades 0-1 and grades 2-3 against SEC^KO^. (**C**) Examples of a fibrinonecrotic focus (dotted outline; note the myocardial necrosis within the outline which is a lighter pink color), (*) a centrally located septic thrombus, and (arrows) intravascular bacteria (deeply basophilic/blue material). Left bar = 100 µm, right bar = 20 µm. (**D**) Examples of an epicardial lesion with saponification (necrosis) of epicardial fat (arrows) and a locally extensive zone of myocardial mineralization (dotted rectangle). Bar = 100 µm. (**E**) Histopathology scoring for the presence or absence of cardiac pathological findings: bacterial thrombi and associated necrosis, intravascular (IV) bacteria, and epicardial fibrin and inflammation.

Most vegetative lesions presented with inflammation adjacent to the vegetations that was almost exclusively heterophilic (neutrophilic) (Fig. 3A). Foci of heterophils infiltrating the myocardium, cellular debris, and necrosis were also observed (Fig. 3A, insets). In the most severe cases (Grade 3), large and coalescing bands of heterophilic infiltrates surrounded the aortic ring (Fig. 3A). Vegetative lesions from *S. aureus* SEC^+^ consistently showed high grade myocardial inflammation that were indistinguishable histologically from those formed by *S. aureus* producing SEC toxoids (SEC_N23A_ or SEC_N23A/F44A/L45A_) (Fig. 3B). Surprisingly, the *S. aureus* SEC^KO^ vegetations that formed on the aortic wall, albeit small, caused multifocal inflammation adjacent to the vegetation (Fig. 3B and S3A). This is in stark contrast to the histopathology from rabbits infected with *S. aureus* SEC^KO^ with no vegetations, which was unremarkable (Fig. S3B). Of interest, septic coronary arterial emboli (coronary arteries containing fibrin or bacterial thrombi) with adjacent myocardial necrosis, and/or intravascular bacteria were observed in rabbits infected with *S. aureus* SEC^+^ and *S. aureus* producing SEC toxoids but absent from those infected with *S. aureus* SEC^KO^ (Fig. 3C and E). Vegetations that penetrated deeper into the pericardium causing epicardial lesions and saponification (necrosis) of epicardial fat was the only other cardiac complication observed in one rabbit infected with *S. aureus* SEC^KO^ (Fig. 3D and E). These observations indicate that 1) intracardial complications associated with embolic events result from the formation of large vegetations produced in the presence of SEC, 2) myocardial inflammation, and its extent, are a direct result of the presence of a vegetation, and 3) these events are largely independent of SEC superantigenicity.

### Superantigenicity differentially contributes to renal and hepatic complications in SAIE

Vegetation fragmentation and metastatic infection occur in one third of SAIE episodes and are associated with hemodynamic and embolic complications in multiple organ systems, including the vascular, pulmonary, gastrointestinal, renal, and hepatic systems (*3*). We had previously observed renal ischemia, infarction, and abscess formation associated with SEC production during SAIE in rabbits (*10*). It remains to be established whether superantigenicity-mediated toxicity significantly contributes to acute kidney and liver injury during SAIE. To address these, experimental rabbits were assessed for kidney and liver lesion pathology. The kidney lesions presented as hemorrhagic, necrotic, or ischemic (Table S3). In the most severe pathology (Grade 3), lesions extended across a large surface of the kidney (Fig. 4A). Kidneys from *S. aureus* SEC^+^ infected rabbits presented with severe pathology in 66% of the animals (Fig. 4B). Similar kidney pathology developed in ~50% of rabbits infected with *S. aureus* producing SEC toxoids (Fig. 4B). Although we did note that Grade 3 pathology was observed only in ~25% of this group, serum levels of blood urea nitrogen (BUN) and creatinine (both biological markers of decreased renal function) were significantly increased in all groups infected with SEC-producing strains compared to SEC^KO^ (Fig. 4C and D).

**Figure 4.**
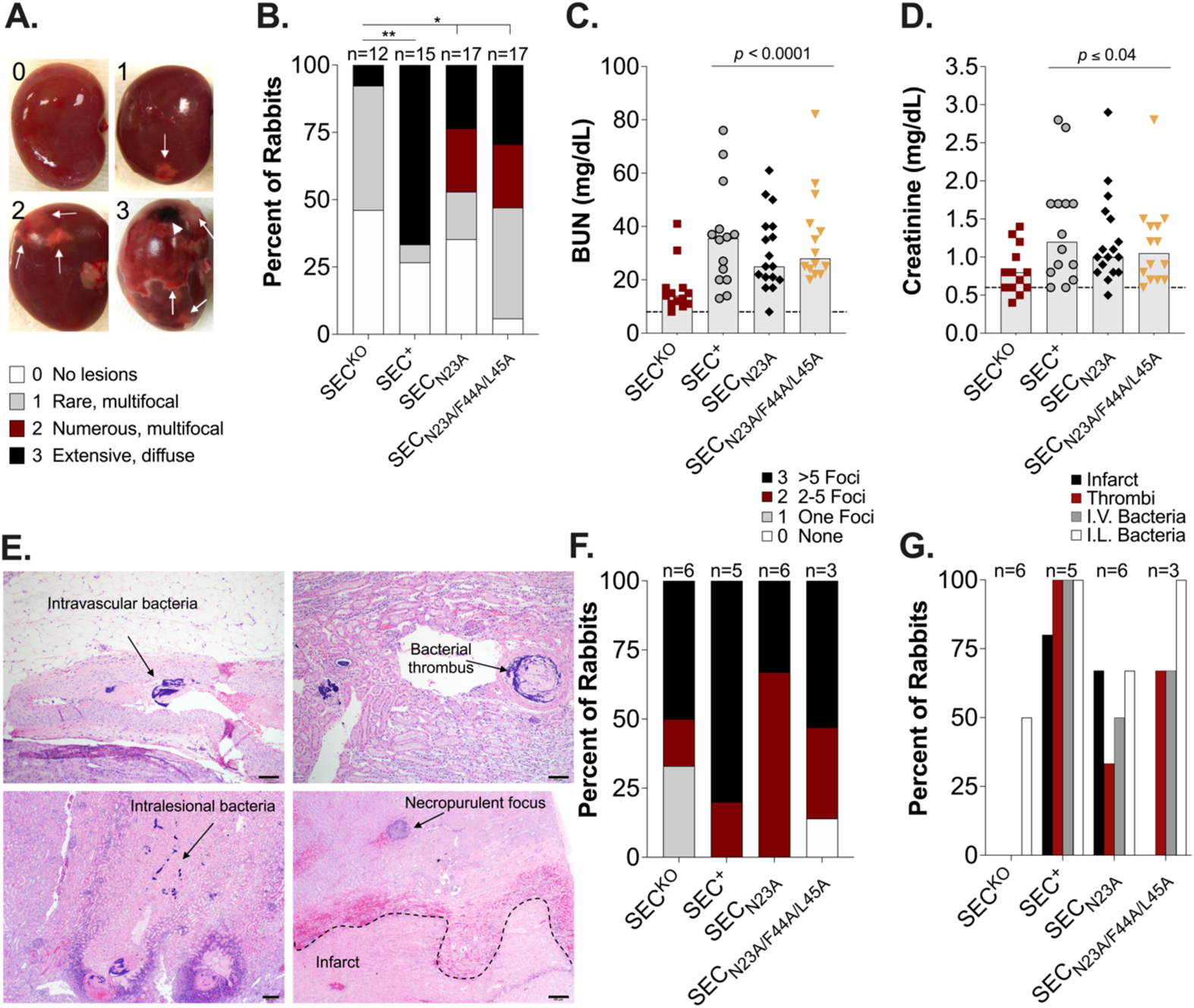
SEC toxoids retain the ability to cause metastatic infections and acute kidney injury. (**A**) Kidney Gross Pathology Grading Scale during rabbit model of native valves infective endocarditis (Grades 0-3). 0 = no lesions, 1 = rare, small (<4mm) multifocal lesions, 2 = numerous large (>5mm) multifocal lesions, 3 = extensive to coalescing to diffuse lesions. Arrows indicate ischemic and/or hemorrhagic lesions, arrowhead indicates a necrotic lesion. (**B**) Scoring of kidney lesions *post-mortem*. *, *p* ≤ 0.0432, **, *p =* 0.0047, Fisher’s exact test between categorical pathology grades 0-1 and grades 2-3 against SEC^KO^. (**C-D**) Serum levels of analytes 48 – 96 h post-infection. Bars represent median value. Dashed line is the average analyte value of all rabbits pre-infection. All groups were tested against pre-infection analyte values and were statistically significant, *p* < 0.03, one-way ANOVA with Holm-Šídák’s multiple comparisons test. (**E**) Examples of histopathologic assessment for the presence or absence of intravascular bacteria (top-left panel, arrow), bacterial thrombi (top-right panel, arrow), intralesional bacteria (bottom-left panel, arrow), and infarcts (bottom-right panel, area below dash line). Bar = 100 µm (top-left and right panels). Bar = 200 µm (bottom-left panel). Bar = 500 µm (bottom-right panel). (**F**) Histopathologic scoring of smaller foci of necropurulent inflammation (Graded 0-3) as shown in **E**, bottom-right panel. 0 = none, 1 = one foci, 2 = 2-5 foci, 3 ≥ 5 foci. (**G**) Histopathologic scoring for the presence or absence of kidney lesions, as shown in **E**: Ischemic necrosis (infarct), bacterial thrombi, intravascular (I.V.) bacteria, and intralesional (I.L.) bacteria.

Histologic analyses of kidney lesions showed that lesions were associated with ischemia and smaller foci of necropurulent inflammation (Fig. 4E). Most lesions from *S. aureus* SEC^+^-infected kidneys contained more than 5 foci of necropurulent inflammation (Fig. 4F), infarctions (areas of ischemic necrosis) with intralesional bacteria in either one or more of these foci of necrosis, intravascular bacteria, and bacterial thrombi (Fig. 4G). Lesions induced by SEC toxoids were indistinguishable histologically from those formed by *S. aureus* SEC^+^ (Fig. 4F). Of interest, lesions from *S. aureus* SEC^KO^-infected kidneys, while exhibiting high grade inflammation (Fig. 4F), failed to exhibit infarction, thrombi, and intravenous bacteria (all signs of embolic events) (Fig. 4G). Hence, SEC-mediated acute kidney injury is largely a result of vegetation formation and fragmentation rather than kidney failure that typically results from toxic shock syndrome (*12*).

Liver pathology presented grossly as pale, streak-shaped ischemic lesions that were focal, multifocal, or wide-spread (Fig. 5A, Table S3). Livers from *S. aureus* SEC^+^ infected rabbits presented with high grade pathology in 80% of the animals (Fig. 5B). Similar liver pathology developed in ~70% of rabbits infected with *S. aureus* producing SEC toxoids (Fig. 5B). Interestingly, rabbits infected with *S. aureus* SEC^+^ had significantly increased mean levels of serum AST (236 U/L) compared to the AST concentration from rabbits infected with *S. aureus* producing SEC toxoids or *S. aureus* SEC^KO^ (AST ≤ 73 U/L) (Fig. 5C). Correspondingly, the ratio of AST to ALT was significantly higher in rabbits infected with *S. aureus* SEC^+^ (Fig. 5D). These increases in liver aminotransferase enzymes are consistent with what is observed in humans during acute liver injury and ischemic hepatitis (*26, 27*). As with the kidneys, ischemic liver lesions were commonly associated with embolic events. Histologic analyses showed ischemic necrosis (well demarcated zones of coagulation necrosis), often surrounded by necropurulent inflammation and/or hemorrhage, and more diffuse centrilobular vacuolation and/or necrosis (indicative of ischemia on a more global scale of the organ) (Fig. 5E and F). It is intriguing that liver lesions from rabbits infected with *S. aureus* SEC^KO^ exhibited high prevalence of infarction and necrosis, when heart and kidney lesions were mostly devoid of embolic complications. It is almost as if the liver was the primary site of infection. Altogether, the data indicate that in addition to the embolic events resulting in localized tissue injury, liver sensitivity to the toxic effects of superantigen activity significantly increases hepatocellular injury.

**Figure 5.**
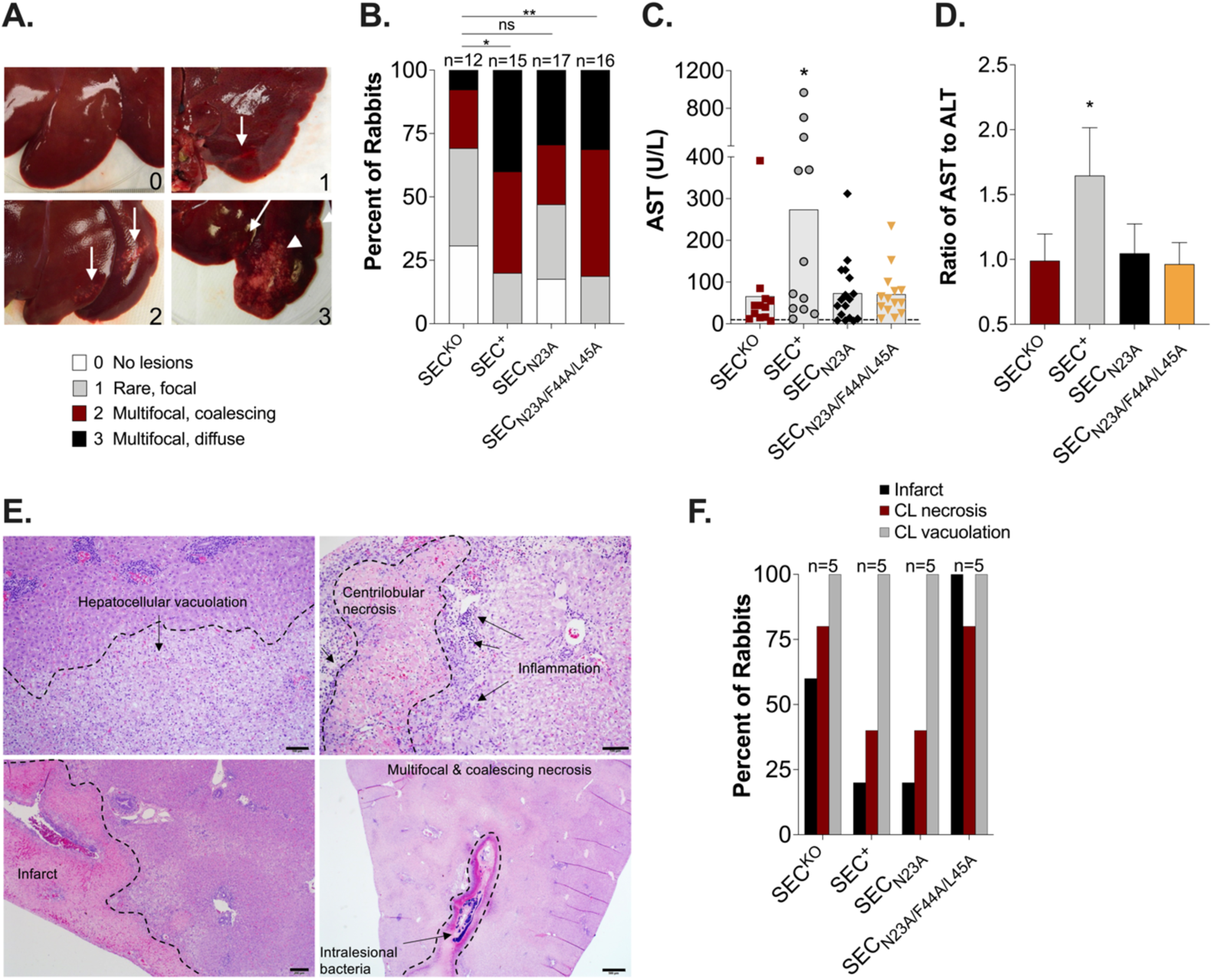
Superantigenicity promotes hepatocellular injury. (**A**) Liver gross pathology grading scale during rabbit model of native valves infective endocarditis (Grades 0-3). 0 = no lesions, 1 = rare, focal streak-shaped lesions, 2 = multifocal to coalescing streak-shaped lesions, 3 = multifocal streak-shaped, extensive to diffuse lesions. Arrows point to streak-shaped ischemic lesions characteristic of grade 1-3, arrowhead indicate wide-spread ischemic lesions characteristic of grade 3. (**B**) Scoring of liver pathology *post-mortem*. *, *p* = 0.02, **, *p* = 0.009, Fisher’s exact test between categorical pathology grades 0-1 and grades 2-3 against SEC^KO^. (**C-D**) Serum levels of analytes 48 – 96 h post infection. Dashed line is the average analyte or ratio value of all rabbits pre-infection. (**C**) Bars represent median value. *, *p* ≤ 0.01, one-way ANOVA with the Holm-Šídák’s multiple comparisons test; *S. aureus* SEC^KO^ compared to other strains. All groups were tested against pre-infection AST values and were statistically significant, *p* < 0.005. (**D**) Data are represented as mean (± SEM). *, *p* ≤ 0.01, Fisher’s exact test comparing ratio values 0-1.8 to values > 1.8 between *S. aureus* SEC^+^ and the rest of the experimental groups. (**E**) Examples of liver lesions observed upon histopathologic assessment. (Top-left panel) locally extensive hepatocellular vacuolation with arrow showing area under dashed line with cytoplasmic vacuolation. (Top-right panel) diffuse centrilobular necrosis (area inside dashed line), surrounded by necropurulent inflammation (arrows). (Bottom-left panel) large zone of necrosis (infarction), area left of the dash line, with multifocal to coalescing portal hepatocellular vacuolation shown on region right of the dash line (arrows show examples). Bar = 100 µm (top-left and right panels). Bar = 200 µm (bottom-left panel). Bar = 500 µm (bottom-right panel). (**F**) Histopathologic scoring for the presence or absence infarcts, centrilobular (CL) necrosis, and centrilobular (CL) vacuolation.

### SEC inhibits neovessel formation in rabbit aortic explants

SEC promoting SAIE and lethal systemic complications largely independent of superantigenicity centers its pathophysiologic effects directly on the endothelium. In acute IE, vegetative lesions develop rapidly with no evidence of repair (*28*). This is followed by fragmentation and septic embolization of cardiac vegetations leading to tissue injury and organ dysfunction, increasing mortality in patients with IE (*29*). Therefore, SEC targeting of endothelial cell function in vascular repair and re-vascularization of injured/ischemic tissues may be central to both localized and systemic pathologies characteristic of SAIE. Angiogenesis is the complex, coordinated process of vascular formation, critical to capillary development and tissue remodeling after injury (*30, 31*). To assess SEC effects on vascular regeneration in a physiologically relevant context, we utilized the *ex vivo* rabbit aortic ring model, which preserves the vessel microenvironment. Ring sections dissected from excised thoracic and abdominal aortas were embedded into a basement membrane matrix to induce angiogenic sprouting along severed edges. Rings were cultured under toxin conditions (SEC_N23A_), antiangiogenic conditions (axitinib), or proangiogenic conditions (basal medium with growth supplements) for 14 days. SEC_N23A_ was used in our *in vitro* studies because: 1) the endothelium does not express TCR, and 2) SEC is a select agent toxin that we can only modify in the SEC_N23A_ background for follow-up studies. Axitinib is an antiangiogenic molecule due to its ability to inhibit signaling via the VEGF-A receptors-1, -2, and -3 thereby preventing VEGF-A overexpression in endothelial cells (*32*). VEGF-A is the most potent pro-angiogenic growth factor (*33, 34*).

In the rabbit aortic ring model of angiogenesis, evidence of microvessel sprouting was observed between days 3-5 in rings maintained in supplemented basal media, with sprouts elongating and branching over 14 days (Fig. 6A, top panels). Of interest, abdominal rings displayed denser microvessel networks with a sprout surface area of 24 ± 1.8 mm^2^ compared to thoracic rings which exhibited a surface area of 15 ± 2.1 mm^2^ (Fig. 6A, top panels). As expected, rings treated with the antiangiogenic molecule axitinib failed to sprout (Fig. S5A). For SEC_N23A_-treated rings, the dose-response analysis showed significant inhibition of sprouting from both thoracic and abdominal aortic rings (Fig. 6B). At the lowest treatment dose of 2.5 μg, SEC_N23A_ significantly decreased the thoracic-ring sprouting area by ~93% over time and the abdominal-ring sprouting area by ~72% (Fig. 6B). Regardless of aortic origin, sprouts that formed on SEC_N23A_ treated rings were detected by day 7 but their further development was stunted (Fig. 6C and D; Fig. S5A and C). These results provide evidence that SEC is an antiangiogenic virulence factor with the potential to effectively inhibit vascular and tissue regeneration by interfering with neovessel formation.

**Figure 6.**
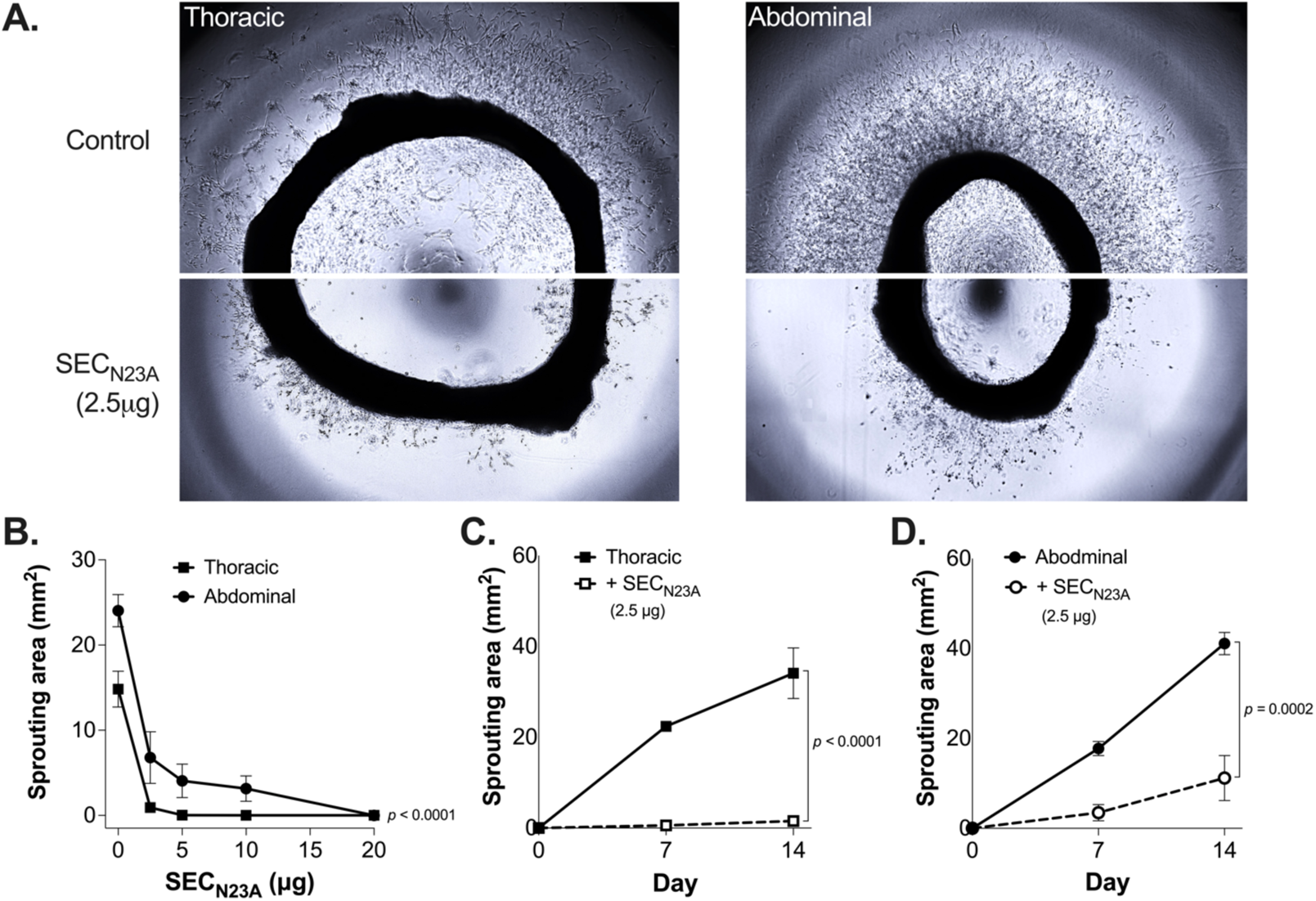
SEC suppresses sprouting of new capillaries in the *ex vivo* rabbit aortic ring model of angiogenesis. Thoracic and abdominal aortic ring explants were cultured in the presence of SEC_N23A_ for 14 days. (**A**) Aortic rings imaged at day 14 showing capillary network formation sprouting from the rings in culture conditions (top panels) versus treatment with 2.5 μg of SEC_N23A_ (bottom panels). (**B**) Quantification of sprouting area (mm^2^) in aortic rings treated with 2.5, 5, 10, 15, and 20 μg of SEC_N23A_ for 14 days. (**C**) Time course analysis of thoracic aortic ring sprouting in the presence (solid squares) or absence (empty squares) of 2.5 μg of SEC_N23A_. (**D**) Time course analysis of abdominal aortic ring sprouting in the presence (solid circles) or absence (empty circles) of 2.5 μg of SEC_N23A_. (**B-D**) Statistical significance was determined by two-way ANOVA with the Holm-Šídák’s multiple comparisons test. P values denote significance between treatment and no-treatment conditions.

### SEC inhibits expression of the proangiogenic factor VEGF-A in human aortic endothelial cells

Re-vascularization and vascular injury repair mechanisms are driven by pro-angiogenic factors released by injured tissues (*30, 31*). Hence, we sought to establish whether SEC modifies the angiogenic response in endothelial cells by altering expression of angiogenesis-related genes. To do so, we profiled the expression of 24 genes important in driving multiple steps of the angiogenic response by RT-qPCR (Table S4). Expression was assessed in immortalized human aortic endothelial cell (iHAEC) at 2, 4, and 6 h under toxin conditions (lipopolysaccharide [LPS] or SEC_N23A_), angiogenesis-enhancing conditions (VEGF-A), or angiogenesis-inhibiting conditions (axitinib). The gene expression profiles were expressed as fold-change from media control, with a cut-off of ≥2-fold or ≤-2-fold as thresholds for 100% increases or decreases in expression levels, respectively. iHAECs grown in standard endothelial cell culture conditions have been shown to produce an angiogenesis protein profile consistent with cells that are triggered to sprout and are therefore already under pro-angiogenic conditions (*35*).

We therefore stimulated iHAECs with bacterial LPS (25 ng/ml) to induce an expression profile characteristic of proinflammatory and pathological angiogenesis (*36*). As expected, LPS-treated cells exhibited a highly pro-inflammatory profile, inducing expression of genes encoding for monocyte chemoattractant protein-1 (MCP-1), granulocyte-monocyte colony stimulating factor-2 (GM-CSF2), and IL-8 (Fig. 7A). The effect of this proinflammatory profile is reflected on the increased gene expression for factors important in extracellular matrix degradation (Serpin E1), growth and survival (Heparin-Binding Epidermal-like Growth Factor [HB-EGF], VEGF-A and VEGF-C), and migration, proliferation and vessel sprouting (pentraxin 3 and angiopoietin-2) (Fig. 7B). VEGF-A stimulation of iHAECs under already pro-angiogenic conditions resulted in induction of its own gene expression while inhibiting genes involved in inflammation, growth, migration, and proliferation (Fig. 7). These results are consistent with an endothelial cell monolayer in homeostasis with no need for further growth. The VEGF-A receptor inhibitor axitinib induced a highly suppressive profile, exhibiting a ≥5-fold downregulation of genes important in ECM degradation (A Disintegrin and Metalloproteinase with Thrombospondin Motifs 1 [ADAMTS-1]), growth and survival (acidic FGF and VEGF-A), proliferation and migration (Tissue Inhibitor of Matrix Metalloproteinase 4 [TIMP-4] and thrombospondin 1), and inflammation (GM-CSF2) (Fig. 7). In stark contrast, iHAECs treatment with SEC_N23A_ at non-toxic concentrations (20 μg/mL; Fig. S6) induces marked inhibition of VEGF-A gene expression, also observed in early time points of axitinib-treated cells (Fig. 7). These results indicate that SEC can exert antiangiogenic effects by inhibiting VEGF-A expression in endothelial cells and help explain SEC_N23A_ suppression of microvessel formation in the aortic ring model.

**Figure 7.**
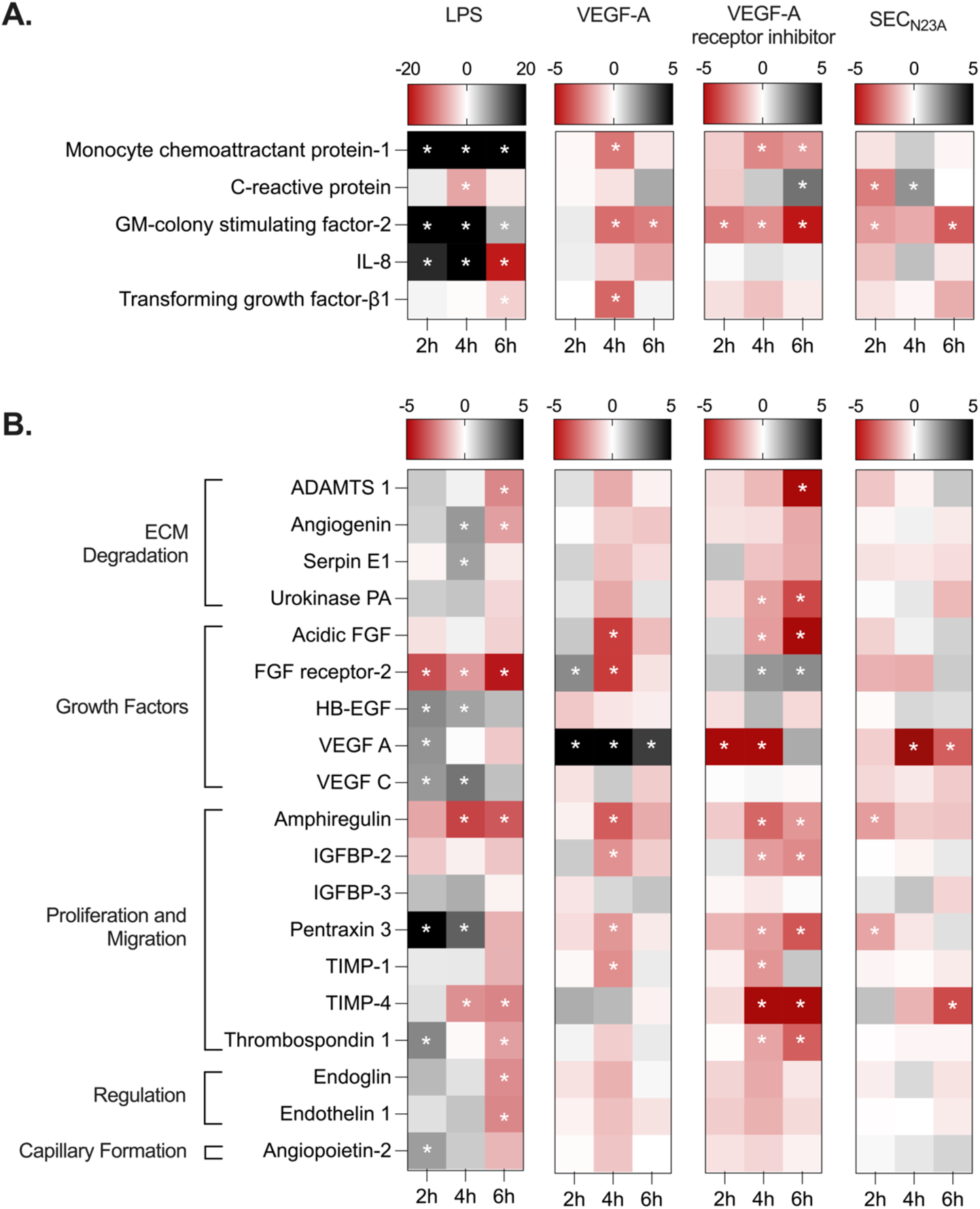
SEC inhibits VEGF-A expression in immortalized human aortic endothelial cells. Expression analysis of 24 angiogenesis-related genes by RT-qPCR in iHAECs treated with lipopolysaccharide (LPS, 25 ng mL^−1^), VEGF-A (20 nM), VEGF-A receptor inhibitor (axitinib, 25 nM), or SEC_N23A_ (20 μg mL^−1^) for 2, 4, and 6 h. (**A**) Gene products that play significant roles in inflammatory responses. (**B**) Gene products that contribute to various angiogenesis processes. *, denotes genes with 2-fold increases or decreases compared to untreated controls.

## Discussion

*S. aureus* SAgs are ubiquitous among human clinical isolates and have been implicated in both colonization and pathogenic mechanisms (*37*). Although their significance as virulence factors has been established, how these multifunctional SAgs specifically contribute to *S. aureus* pathogenesis has become a pressing question (*38–40*). Current evidence indicates that the SAgs SEC, TSST1 and select *egc* toxins play a novel and essential role in the etiology of SAIE by yet uncharacterized mechanisms (*10, 11*). Still, in SAg-mediated illnesses, superantigenicity is placed as a triggering event mediating or exacerbating pathological responses during *S. aureus* infection. It has been proposed, albeit not directly tested, that superantigenicity leading to hypotension and immune dysregulation allows for bacterial immune evasion and persistence, while direct interaction with the heart endothelium promotes endothelium dysfunction and disease progression (*10, 11, 38*). With the use of *S. aureus* producing SEC inactivated in MHC-II and/or TCR binding, we provide evidence that superantigenicity is not the primary mechanism by which SEC promotes vegetation formation, cardiac toxicity, and extracardiac complications such as acute kidney injury. Our results are consistent with published studies noting that not all SAgs promote SAIE to the same extent (*11*). SEG and SEN have comparable T cell mitogenic activity as SEA, SEB, and TSST1 (*41, 42*), yet they exhibit decreased efficiency in promoting SAIE. Our studies highlight the critical contribution of SAgs in life-threatening pathologies by at least two novel mechanisms: 1) interference with vascular injury repair, and 2) inhibition of re-vascularization of injured or ischemic tissues.

During SAIE, vascular injury results from mechanical damage and/or toxin-mediated damage of the endothelium. Re-endothelialization of injured vessels restores their integrity and is essential to limit the progressive loss of the endothelial layer and continual deposition of host factors (e.g. platelets, fibrin, leukocytes) that leads to mural thrombus formation (*37*). In the case of SAIE, re-endothelialization defects would promote expansion of the vegetative lesion and growth of the vegetation. We have previously shown that the SAg TSST1 directly causes dysregulated activation of iHAECs and disrupts re-endothelialization in an *in vitro* wound-healing assay, suggesting a role of SAgs in endothelial dysfunction and a possible explanation for their role in SAIE (*38*). Both re-endothelialization and sprout elongation are a function of cell proliferation and migration, and their impairment results in angiogenesis defects (*43*). Angiogenesis is a highly complex multistep process requiring endothelial cell activation, basement membrane degradation, sprout formation and elongation, and capillary network stabilization (*30, 31*). These processes are also supported by endothelium-associated cells, such as pericytes, fibroblasts, monocytes, and smooth muscle cells (*44, 45*). These cell types are not present in standard monoculture *in vitro* wound healing assays. For the study of angiogenesis, the *ex-vivo* rabbit aortic ring model is a more physiologically relevant model (*30, 46*). With this model, we provide conclusive evidence that SEC disturbs branching microvessel formation, effectively inhibiting angiogenesis.

We propose that SEC inhibition of angiogenesis occurs via a mechanism dependent on down-regulation of one of the most potent inducers of angiogenesis, VEGF-A (*33, 34, 47, 48*). VEGF-A is a proangiogenic growth factor produced in virtually all vascularized tissues (*33, 48*). It promotes endothelial cell survival and mitogenic stimuli (*49, 50*), re-endothelialization of denuded arterial walls to restore the integrity of the endothelium (*51*), and growth of new capillaries to restore perfusion to ischemic tissues (*52, 53*). Physiological levels of VEGF-A maintain vascular homeostasis and are vasculoprotective, while VEGF-A up-regulation is critical to trigger sprouting angiogenesis and wound repair mechanisms (*54–56*). We provide evidence that SEC inhibits expression of *VEGFA* in iHAECs. This inhibition peaks between 2 – 6 hours after SEC exposure. Similar inhibition is observed in cells exposed to axitinib, a potent inhibitor of the VEGF-A receptor (*32*). Inhibition of the VEGF-A signaling pathway, regardless of the mechanism (e.g. VEGF-A neutralization or VEGF-receptor inhibition), results in endothelial dysfunction and vascular toxicity, including thrombosis, inappropriate pruning of the arterial vasculature, and re-vascularization defects during ischemic injury (*57–59*). These events can help explain the pathophysiology typical of SAIE.

SAIE has a high mortality rate owing to the high incidence of both intracardiac complications arising from the rapid local spread of the infection and the high incidence of embolization of septic vegetation fragments (*1, 3, 60*). Lodging of septic emboli within terminal blood vessels causes localized ischemia and infarction in multiple organ systems. In humans and in experimental rabbits, these complications can be manifested as myocardial infarction, kidney and/or liver injury, and strokes (*60*). Of these, kidney injury leading to acute renal insufficiency with progression to acute renal failure is tightly associated with SAIE severity, development of septic shock, and IE lethality (*7*). Our study demonstrates that SEC-associated kidney injury is a consequence of its role in vegetation formation and embolization of cardiac vegetations. The kidney lesions observed upon examination ranged from ischemic or hemorrhagic to necrotic. In this context, VEGF-A deficits and angiogenesis defects in general can lead to persistent vascular injury and insufficient re-vascularization, overall impairing the wound healing process allowing for renal deterioration and bacterial growth. The liver has central roles in clearing circulating bacteria and their toxins and is key in initiating or amplifying inflammatory responses during systemic infections (*7*). Yet, literature on the effects of SAIE on liver injury is scarce (*3*). In our study, increases in serum aminotransferases correlates with increases in IL-6 and LDH in rabbits infected with *S. aureus* SEC^+^. Therefore, in contrast to the kidneys, in the liver, superantigenicity significantly contributes to cellular injury.

Our current understanding of SAg involvement in the pathogenesis of many life-threatening *S. aureus* infections such as bacteremia, pneumonia, and IE has been tied to complications arising from the extreme effects of superantigen activity. While adaptive immune system activation is the hallmark of staphylococcal SAgs, we provide evidence that this is not their only biological function involved in promoting life-threatening disease pathologies. It is clear that SAgs do much more than previously anticipated or expected. Therefore, it becomes fundamental to further understand the involvement of superantigenicity-independent mechanisms in other invasive and life-threatening diseases as well.

Recently, the SAg SE*l*-X was found to inhibit neutrophil function via a sialic acid-binding motif uniquely present in this SAg (*40*). Meanwhile, TSST1 was reported to induce activation of epithelial cells through a dodecapeptide close to the base of the central α-helix of the molecule (*39*). The dodecapeptide sequence is found in all staphylococcal SAgs, yet its effects on non-hematopoietic cells is poorly characterized (*12, 39, 61*). The relevance of the SEC dodecapeptide in endothelial cell function and *S. aureus* diseases such as IE is not known but it is at the center of current studies.

Altogether, our study demonstrates that superantigenicity independent effects of SEC are essential to the pathogenesis and pathophysiology of SAIE. The data further shows that SEC is an anti-angiogenic toxin that inhibits sprouting of new capillaries from the activated endothelium, which would effectively impair re-vascularization of injured tissues. It is becoming evident that SAgs like TSST1 and SEC promote IE by interfering with vascular repair and the angiogenic process. One of the mechanisms by which SEC may exert its anti-angiogenic effects is through the inhibition of endothelial cell expression of one of the key inducers of angiogenesis, VEGF-A. If so, deficient angiogenesis may be at the center of many pathologies associated with SAg-producing *S. aureus* infections. Given the reach of the vascular endothelium into every organ system and the prevalence of SAgs among both methicillin-susceptible and resistant *S. aureus* strains, SAgs have the potential to promote pathologies in every organ that they encounter.

## Materials and methods

### Bacterial strains and growth conditions

Community-associated, methicillin-resistant USA400 strain MW2 was originally obtained in the Upper Midwest from a young patient who succumbed to necrotizing pneumonia. *S. aureus* MW2 encodes for the SAgs SEC, SEA, SEH, SEK, SEL, SEQ, and SE-*like* X. SEC is produced at 60-100 μg/ml in liquid culture, while the rest are produced at 0.0001-0.03μg/ml. Staphylococcal strains were used from low-passage-number stocks. All staphylococcal strains were grown in Bacto™ Todd Hewitt (TH) (Becton Dickinson) broth at 37°C with aeration (225 rpm) unless otherwise stated. Strains and plasmids used in this study are listed in Table S1. Plasmids used for complementation were maintained using carbenicillin (100 µg/ml) in *E. coli* DH5⍺. For endocarditis experiments, strains were grown overnight, diluted, and washed in phosphate buffered saline (PBS - 2mM NaH_2_PO_4_, 5.7 mM Na_2_HPO_4_, 0.1 M NaCl, pH 7.4).

### Selection of mutations inactivating the SEC TCR binding site (N23A) or the MHC-II binding site (F44A/L45A)

Asn^23^ is a highly-conserved, surface exposed residue located in a cleft between the O/B fold and β-grasp domain of SAgs (*62*). It forms hydrogen bonds with the backbone atoms of the complementarity-determining region (CDR) 2 of the Vβ-TCR (*63*). As such, Asn^23^ contact with the Vβ-TCR has one of the greatest energetic contributions of the complex. Mutations in this position, such as N23A or N23S, greatly destabilize the Vβ-TCR:SAg interaction with profound effects in SAg activity (*63*). SEC_N23A_ has no detectable binding to Vβ-TCR as measured by surface plasmon resonance, no proliferative T-cell responses in thymidine-incorporation assays at concentrations up to 30 μg/ml (*63*), no lethality or signs of TSS in rabbits vaccinated subcutaneously with 25 μg three times every two weeks (*16*), and no lethality in rabbits after intravenous injection at 3,000 μg/kg (*15*). Phe^44^ and Leu^45^, conserved among all enterotoxins, are located on a protruding hydrophobic loop that directly contacts MHC-II and forms strong electrostatic interactions with the α-chain (*20, 64*). Leu^45^ is the most extensively buried amino-acid residue in the SEC:MHC-II interface, but mutations in either residue (Phe^44^ or Leu^45^) effectively inactivate SEC binding (*18–21, 64*). F44S alone is 1000-fold less efficient in MHC-II binding resulting in a concomitant reduction of IL-2 in T-cell proliferation assays (*19, 65*).

### Construction of chromosomally complemented toxoid strains

SEC is a CDC designated select agent. As such, we are not allowed to use the wildtype copy of the gene in recombinant studies. For this reason, all PCR products generated in the making of the toxoid complement strains either included the permissible N23A mutation or was only a partial amplification of *sec* missing either the TCR or MHC-II domain. Each step of plasmid construction was verified by Sanger sequencing to contain the N23A TCR mutation. PCR amplification was performed using Phusion polymerase (New England Biolabs; NEB) unless otherwise noted. *S. aureus* expressing SEC_N23A_ or SEC_N23A/F44A/L45A_ were made by markerless chromosomal complementation in MW2Δ*sec* with the genes expressed under the control of the native promoter and terminator (*66*). The SEC_N23A_ gene sequence was created by amplifying two fragments from MW2 with primer sets pJB38xN23ApromF/promN23R and termN23F/pJB38xN23AtermR. The chromosomal complementation plasmid, pJB38-NWMN29-30 (RRID:Addgene_84457), was digested with EcoRV and PCR products inserted by Gibson Assembly (NEB) as previously described, creating pKK29 (*67*). SEC_N23A_ was amplified from pKK29 with the primer set pUC19SECN23AptF/pUC19SECN23AptR and inserted by Gibson Assembly into pUC19 (RRID:Addgene_50005) linearized with KpnI and EcoRI, to create pKK33. The MHC-II binding site mutations were introduced into pKK33 by site-directed mutagenesis (QuickChange II, Agilent Technologies) using the primer set SECF44A/L45Afor/ SECF44A/L45Arev, creating pKK39. The SEC_N23A/F44A/L45A_ gene sequence was amplified from pKK39 with primer set pJB38xN23ApromF/ pJB38xN23AtermR and inserted into pJB38-NWMN29-30 as described above, creating pKK42. pKK29 and pKK42 were electroporated into *S. aureus* RN4220 and moved into MW2Δ*sec* by generalized transduction with ϕ11(*68*). *S. aureus* strains containing plasmids were selected for with chloramphenicol (20 µg/ml) at 30°C. Allelic exchange was performed as previously described (*66*), chromosomal insertions detected by PCR with primer set XNWMN2930F/XNWMN2930R, and verified by Sanger sequencing. Primers were purchased from Integrated DNA Technologies (Table S2).

### SEC purification and T cell proliferation assay

SEC was purified from *S. aureus* strain FRI913 in its native form by ethanol precipitation and thin-layer isoelectric focusing as previously described (*69*). Preparation of SEC resulted in a single band by Coomassie blue stain. Toxin preparations were tested for lipopolysaccharide (LPS) contamination with the Toxin Sensor Chromogenic LAL Endotoxin Assay following the manufacturer’s instructions (GenScript). SEC preparations had < 0.02 ng of LPS per 20 µg of SEC^+^. For analysis of superantigen activity in *S. aureus* expressing SEC_N23A_, SEC_N23A/F44A/L45A_, SEC, or SEC^KO^, cell-free supernates were collected from bacteria grown overnight, centrifuged, and filtered (0.22-µm pore size, MilliporeSigma). Mononuclear cells were isolated from human peripheral blood samples (PBMC) originating from 3 donors by density gradient centrifugation using ficoll-paque PLUS (UW–Madison Minimal Risk IRB Protocol 2018-0304-CR004). PBMCs were labelled with 5µM Cell Trace Violet (CTV) dye (Thermo Fisher Scientific) following the manufacturer’s protocol. CTV labelled cells were plated in 96-well plates at a concentration of 2 × 10^4^ cells/well in RPMI 1640 medium supplemented with 10% heat-inactivated fetal bovine serum, 5% heat inactivated bovine calf serum, 3% human AB serum, 1% L-glutamine and 1% penicillin/streptomycin. Purified SEC (10μg), or cell-free supernates at 0.3, 2, or 10-μl volumes were added to the wells. After 6 days, cell division of CD4^+^ T cells was assessed flow cytometrically by measuring dilution of CTV fluorescence intensity. Proliferation response was quantified using the FlowJo proliferation platform analysis tool and data presented as replication index or percent of CD4^+^ T cell divided.

### Rabbit model of SAIE

The rabbit model of IE was performed as previously described with some modifications (*10*). 2-3 kg New Zealand White Rabbits were obtained from Bakkom Rabbitry (Red Wing, MN) and anesthetized with ketamine (dose range: 10-50 mg/kg) and xylazine (dose range: 2.5-10 mg/kg). Mechanical damage to the aortic valve was done by introducing a hard, plastic catheter via the left carotid artery, left to pulse against the valve for 2 h, removed, and the incision closed. Rabbits were inoculated via the marginal ear vein with 2 x 10^7^-4 x 10^7^ total CFU in PBS and monitored 4 times daily for a period of 4 days. For pain management, rabbits received buprenorphine (dose range: 0.01 – 0.05 mg/kg) twice daily. At the conclusion of each experiment, bacterial counts were obtained from heparinized blood (50 USP units/mL). Rabbits were euthanized with Euthasol (Virbac) and necropsies performed to assess overall health. Spleens were weighed and used as an infection control, kidney and liver gross pathology was graded using gross lesion pathology scale (Table S3), aortic valves were exposed to assess vegetation growth, and vegetations that formed were excised, weighed, and suspended in PBS for CFU counts. A minimum of 4 rabbit hearts from each infection group were placed in 10% neutral buffered formalin and further processed by the Comparative Pathology Laboratory at the University of Iowa for histopathological analyses. Vegetation weight and bacterial counts cannot be obtained from hearts prepared for histology. Experiments were performed according to established guidelines and the protocol approved by the University of Iowa Institutional Animal Care and Use Committee (Protocol 6121907). Number of rabbits per infection group were as follow: *S. aureus* SEC^+^ (n=12), *S. aureus* SEC^KO^ (n=15), *S. aureus* SEC_N23A_ (n=17), and *S. aureus* SEC_N23A/F44A/L45A_ (n=17). Data from rabbit experimental groups are a result of at least 3 independent experiments.

### Serum analysis

Rabbit serum was obtained from heparinized blood (50 USP units/mL) collected before infection and at 48 h, 72 h, and 96 h post infection. Blood was centrifuged at room temperature at 5000 x g for 10 min. The collected supernatant was centrifuged for an additional 5 min, filter sterilized using a 0.2 μm filter, and stored at −80°C for further analysis. Serum samples were sent to the University of Iowa Diagnostic Laboratories and evaluated for the following serum analytes: aspartate aminotransferase (AST; U/L), alanine aminotransferase (ALT; U/L), blood urea nitrogen (BUN; mg/dl), creatinine (mg/dl), and lactate dehydrogenase (LDH; mg/dl). IL-6 was quantified from serum samples using the R&D Systems DuoSet Rabbit IL-6 ELISA kit according to manufacturer’s instructions. Serum samples were diluted 1:10 in reagent diluent prior to use. The optical density (O.D.) was determined using a TECAN M200 plate reader (Tecan Group Ltd.) set to 450 nm with wavelength correction set to 540 nm. A standard curve was created by linear regression analysis of the IL-6 concentration versus O.D., log-transformed (GraphPad Prism 8).

### Histopathology scoring

Fixed tissues were routinely processed, cut at 5 µm, and hematoxylin and eosin (H&E) stained or Gram stained. The slides were scored in a semi-blinded fashion, with the pathologist only made aware of control animals. Histopathologic scoring was developed specifically for this animal model, as outlined in Gibson-Corley et. al, 2013 (*70*) and explained in Table S3. Rabbits infected with *S. aureus* SEC^KO^ with no vegetations did not exhibit pathology at the end of experimentation (n=3). Hence, all hearts of *S. aureus* SEC^KO^ infected rabbits with visible vegetations were processed (mean size 2.7 ± 1 mm^2^, n=4). Heart, kidney and liver sections from rabbits infected with *S. aureus* SEC^+^, *S. aureus* SEC_N23A_, and *S. aureus* SEC_N23A/F44A/L45A_ were selected randomly based on presence of vegetations (mean size 6.6 ± 2 mm^2^, n=15) and gross pathology lesions in the kidney and liver.

### SEC*_N23A_* expression and purification

SEC_N23A_ (C-terminus 6xHis-tagged) was expressed in *E. coli* BL21 (DE3) Star strain harboring pET25bHSVdelTEV. Following ethanol precipitation, toxins were purified through Ni^2+^-affinity chromatography using a HiTrap Heparin HP column (GE Healthcare). Protein capture was verified via visualization of a single band on SDS-PAGE following Coomassie blue staining. Detoxi-Gel endotoxin removing resin (Thermo Scientific) was used to reduce LPS contamination to <0.001ng of LPS per 30μg of toxin was obtained. Endotoxin levels were quantified using ToxinSensor™ Chromogenic LAL Endotoxin kit (GenScript) to ensure toxin purity.

### *Ex-vivo* rabbit aortic ring model of angiogenesis

Mixed-sex New Zealand white rabbits, 2-3 kg, were purchased from Charles River Laboratories (Massachusetts) and maintained at Charmany Instructional Facility at the School of Veterinary Medicine (SVM) of the University of Wisconsin–Madison, according to established guidelines and the protocol approved by the Institutional Animal Care and Use Committee (Protocol V006222-A02). Aortic ring explants were cultured by modifying the thin-layer method (*71*). Briefly, the thoracic and abdominal aortas were excised immediately after euthanasia, cleaned to remove excess fascia and connective tissue, dissected into 1 – 1.5 mm cross-sections, and embedded into GFR-Matrigel coated wells (200 μL phenol red-free Corning^®^ Matrigel^®^ Growth Factor Reduced (GFR) Basement Membrane Matrix) in a 48-well plate format. After 10 min polymerization at 37°C, 5% CO_2_, 200-μL supplemented Medium 200 was added and plates incubated at 37°C, 5% CO_2_ for up to 14 days. Medium 200 contained LSGS (see Human aortic endothelial cell culture and RNA extraction), 100 U mL^−1^ penicillin-streptomycin, 2.5 μg mL^−1^ amphotericin B, and designated treatment. Media (± treatments) was changed every 3-5 days. Rings were treated with SEC_N23A_ at 2.5, 5, 10, and 20 μg and axitinib at 10 μM. Rings treated initially with 20 μg were switched to 10 μg following the first media change to maintain toxin pressure while limiting potential cytotoxic effects. Equal numbers of thoracic and abdominal aortic rings were used for each treatment condition as follows: media control (n=24), axitinib (n=8), SEC_N23A_ 2.5 μg (n=6), SEC_N23A_ 5 μg (n=9), SEC_N23A_ 10 μg (n=9), and SEC_N23A_ 10 μg (n=8). All treatment conditions were assessed from 3 separate rabbits with the exception of SEC_N23A_ at 2.5 μg, which was completed in 2 rabbits. Phase-contrast images were captured on days 0, 7, and 14 with the Leica DMi8 equipped with a Tokai Hit stage-top incubator set to 37°C, 5% CO_2_ using a HC PL FLUOTAR 4x/0.13 objective lens. Sprouting was assessed using ImageJ (*72*).

### Human aortic endothelial cell culture and RNA extraction

Immortalized human aortic endothelial cells (iHAECs) are a recently established cell line shown to retain phenotypic and functional characteristics of primary cells (*38*). iHAECs were cultured at 37°C, 5% CO_2_ in phenol red–free, endothelial cell basal medium (Medium 200) supplemented with low-serum growth supplement (LSGS, final concentrations of: FBS 2%, hydrocortisone 1μg mL^−1^, human epidermal growth factor 10 ng mL^−1^, basic fibroblast growth factor, 3 ng mL^−1^, heparin 10 μg mL^−1^), both from Gibco Life Technologies), as previously described (*38*). LSGS contains the final concentrations of: FBS 2%, hydrocortisone 1μg mL^−1^, human epidermal growth factor 10 ng mL^−1^, basic fibroblast growth factor, 3 ng mL^−1^, heparin 10 μg mL^−1^). All experiments were conducted with iHAECs at passages 4-10 from a single clone seeded at 5 x 10^5^/well in a 6-well plate. iHAECs were treated with LPS (25 ng mL^−1^), VEGF-A (20 nM), axitinib (25 nM), or SEC (20 μg mL^−1^) for 2, 4, and 6 h. Data are the result of three biological replicates. RNA was extracted at the indicated time points using the miRNeasy Mini Kit (Qiagen), according to the manufacturer’s instructions, and stored at –80°C. Contaminating genomic DNA (gDNA) was removed using the Turbo DNA-Free™ Kit (Invitrogen). RNA quantity and purity were assessed using the NanoDrop™ 1000 (Thermo Fisher) with median A260/280 values of 1.93 and A260/230 of 1.55.

### cDNA synthesis and real-time reverse-transcription PCR

cDNA synthesis was carried out using DNA-free RNA with the RT First Strand Kit according to manufacturer’s instructions (Qiagen). Each cDNA sample was used at 120 ng/well in 96-well plates. qPCR was conducted using the PowerUP SYBR Green Master Mix (ThermoFisher Scientific, USA) with each reaction having 5 μL of PowerUP SYBR Green Master Mix, 2 μL of 1 μM forward and reverse gene-specific primers (Table S4), and 3 µL of diluted cDNA experimental sample in a final reaction volume of 10 µL. Reactions were run with the following cycle profile: initial denaturation −95°C 10 minutes, 40 cycle amplification/elongation −95°C 15 seconds followed by 60°C 1 minute. gDNA contamination from each sample was assessed using a total of 120 ng of RNA (no reverse transcriptase) with reaction conditions as listed above with all samples having C_T_ values >35. No-template controls were run with each qPCR plate and had C_T_ values >38. All qPCR reactions were run and analyzed on the QuantStudio 3 Real-Time PCR System with analysis software v1.5.1 (Applied Biosystems). Each timepoint was collected in biological triplicates with qPCR reactions from each experimental sample done in technical triplicates.

### Statistical analyses

The log-rank, Mantel Cox test was used for statistical significance of survival curves. Normality was assessed using the D’Agostino & Pearson test along with associated Q-Q plots for data distribution. For comparison across means log-transformed data was used and statistical significance was determined by using one-way analysis of variance (ANOVA) with Holm-Šídák’s multiple comparisons test for the following data sets: vegetation size, vegetation CFU, blood CFU, spleen size, BUN, creatinine, AST, ALT, LDH, and IL-6. Statistical significance for gross pathology data was determined using Fisher’s exact test along with calculated odds ratios and 95% confidence intervals. Statistical significance for sprouting area in the aortic ring model was determined by two-way ANOVA with Holm-Šídák’s multiple comparisons test. Statistical significance of virulence factor production was determined with the nonparametric Kruskal-Wallis test. α = 0.05 (GraphPad Prism 8).

## Supporting information

supplementary materials for the preprint site

## Acknowledgements

We thank the University of Iowa Comparative Pathology Laboratory and University of Iowa Diagnostic Laboratories for histology and serum analysis services.

## Funding

This work was supported by National Institutes of Health (NIH) grant R01AI34692-01 to W.S-P, NIH grant 5T32AI007511-23 to P.M.T., and NIH grant T32GM008365 to K.J.K.

## Author Contributions

K.J.K., S.S.T., J.E.G. and W.S-P conceptualized and designed experiments, analyzed the data, and wrote the manuscript. K.J.K, P.M.T, A.N.F, K.K, and W.S-P carried out *in vivo* rabbit experiments. K.N.G-C provided intellectual and technical support on histopathological analysis and gross pathological grading. S.S.T. carried out ex vivo experiments. X-J. W carried out qPCR analyses. N.S.B. carried out T cell proliferation assays, analyzed the data. All authors reviewed the manuscript.

## Competing Interests

K.J.K. is currently an employee of Integrated DNA Technologies, which sells reagents used or like those used in this manuscript.

## Data and materials availability

All the data needed to evaluate the conclusions in this manuscript are present in the manuscript and/or supplementary materials.

## Supplementary materials and methods

### SDS-PAGE and quantitative western blotting

*S. aureus* strains were grown for 24 h in TH. 5 mL of culture was precipitated in 20 ml of 100% EtOH (Final Concentration 80% EtOH) and stored at 4°C overnight. Precipitates were collected by centrifugation at 4°C for 10 min at 1000 x g. Supernatants were poured off and precipitates were allowed to dry, resuspended in 250 µl of PBS, and centrifuged at room temperature for 10 min at 21,000 x g. The supernatants were collected and total protein quantified by Bradford assay according to manufacturer’s instructions using bovine serum albumin as a standard (Bio-Rad Laboratories). Samples were normalized to a protein concentration of 0.4 mL^−1^, diluted 1:20 in PBS, mixed with 2X sample buffer, boiled for 5 min, and electrophoresed along with purified SEC as a standard curve on a 12% Mini-PROTEAN^®^ TGX Stain-Free™ gel (Bio-Rad Laboratories) at 200V for 40 min. Samples were transferred to an Immobilon-FL polyvinylidene fluoride (PVDF) membrane (MilliporeSigma) by electroblotting at 100V, 250mA for 90 min at 4°C in Towbin Buffer with 20% EtOH substituted for methanol. Membranes were blocked for 1 h at room temperature with 10% nonfat dry milk and 5% heat inactivated horse serum (Gibco Life Technologies) in PBS and then incubated overnight at 4°C with rabbit polyclonal anti-SEC antibody (1:10,000) in 5% nonfat dry milk and 2.5% heat inactivated horse serum in PBS. The membrane was washed with PBST (PBS with 0.1% Tween 20) and incubated with goat anti-rabbit antibody IRDye 800CW (LI-COR, 1:20,000) for 1 h at room temperature. Blots were washed with PBST and visualized using an Odyssey CLx (169 µm resolution, auto intensity 800 nm channel) and analyzed with Image Studio software (LI-COR). SEC production was quantified by standard curve method using purified SEC and densitometrical analysis. Data is represented from 5 independent experiments with technical triplicates.

### *S. aureus* growth curves

Overnight cultures were diluted to an OD_600_ 1.0 in TH. 10 µl of diluted culture was added to 190 µl of TH in a 96-well round, clear bottom plate in technical triplicate (Corning). Plates were sealed using Breathe-Easy sealing membrane (Diversified Biotech). Growth was measured using a TECAN M200 plate reader overnight with the following settings: 37°C, 30 min kinetic interval, orbital shaking: amplitude 3 mm for 10 s before reads, 1 read/well at 600 nM, 25 flashes, 9 nM bandwidth, 100ms settle time, and 27 min wait time between reads with orbital shaking: amplitude 3mm for 26.5 min. Data is represented by three independent experiments.

### Erythrocyte lysis assays for hemolysin production

Erythrocyte Lysis assays were performed as previously described (*73*). Overnight cultures were diluted to an OD_600_ 1.0 in PBS and 5 µl spotted onto TSA II agar plates with either 5% rabbit or sheep blood (Becton Dickinson). Plates were incubated for 24 h at 37°C with 5% CO_2_. Zones of hemolysis were measured and quantified using ImageJ. Data is represented by three independent experiments done with technical duplicates.

### MTS assay

An MTS assay was used to determine cell viability. Cells were seeded at 7,000 cells/well into gelatin-coated 96-well plates and grown overnight to near confluency. Media was removed and replaced with 100 μL of media containing increasing concentrations of SEC or SEC_N23A_ at 20 μg mL^−1^ followed by overnight incubation. 20 μL of CellTiter 96® AQueous One Solution was added to each well followed by a 1 h incubation at 37°C, 5% CO_2_. A plate reader was used to read absorbance at 490 nm.

Three biological replicates were conducted with each containing three technical replicates. The data were normalized so that untreated cells were considered 100% activity by dividing the absorbance of treated cells by the absorbance of untreated cells.

## Supplementary figures

**Figure S1.**
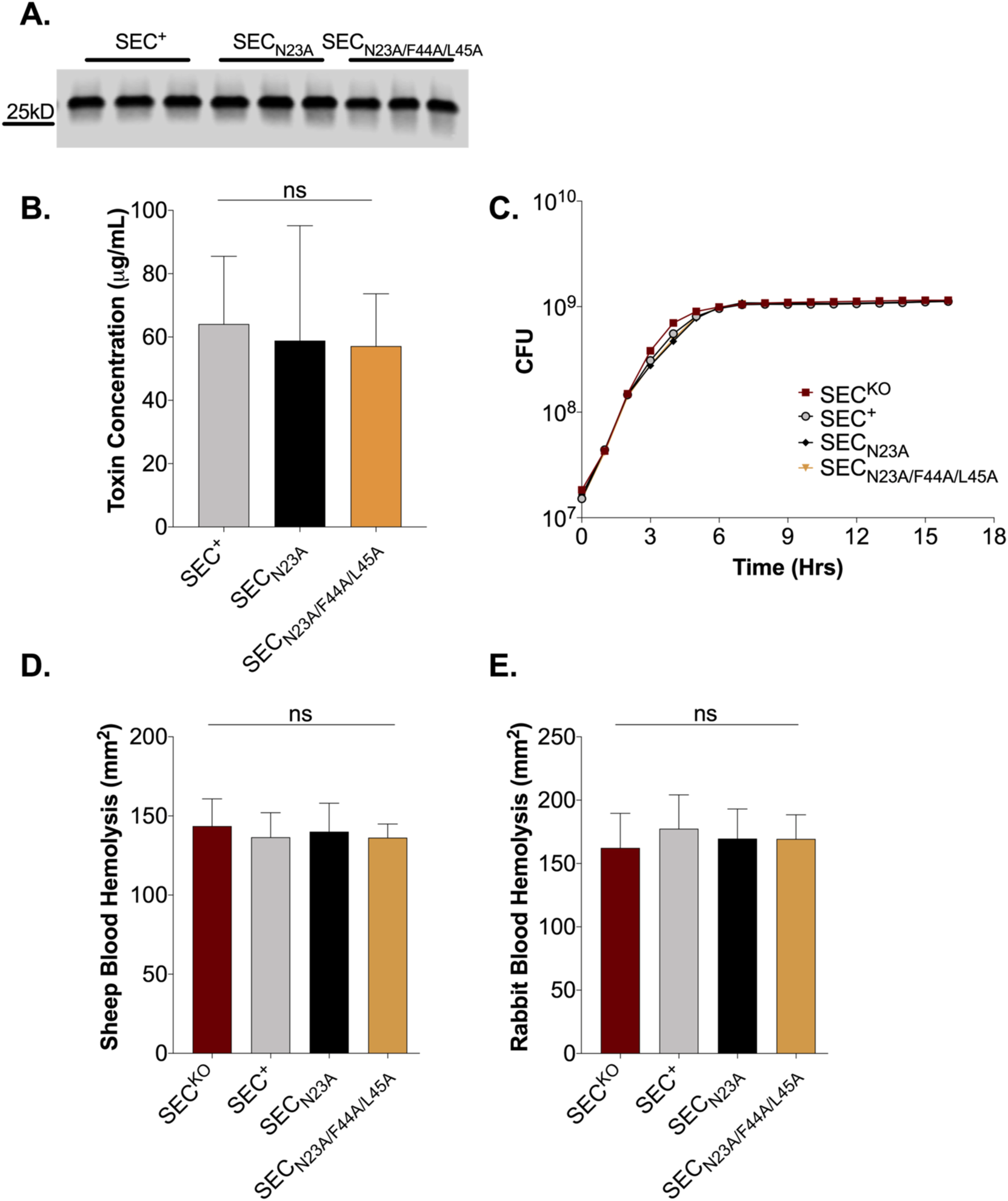
Virulence factor production and growth of strains producing SEC binding site inactivated toxoids. (**A**) Representative western blot images of SEC production in wild-type and SEC complement strains grown overnight in Todd Hewitt (TH) medium shown in technical triplicate. (**B**) Quantitative western blot analysis of SEC production. (**C**) Growth curve of *S. aureus* SEC^KO^, SEC^+^, SEC_N23A_, SEC_N23A/F44A/L45A_ grown overnight in TH. (**D**) Relative levels of hemolysin production as measured in a sheep erythrocyte lysis assay. (**E**) Relative levels of ⍺-hemolysin production as measured in a rabbit erythrocyte lysis assay. (**D, E**) Overnight cultures of *S. aureus* were washed and spotted onto blood agar plates. Zones of hemolysis were measured after overnight growth. Data are represented as mean (± SD). Statistical significance was determined by one-way ANOVA and nonparametric Kruskal-Wallis test. *p* values ≤ 0.05 are considered statistically significant.

**Figure S2.**
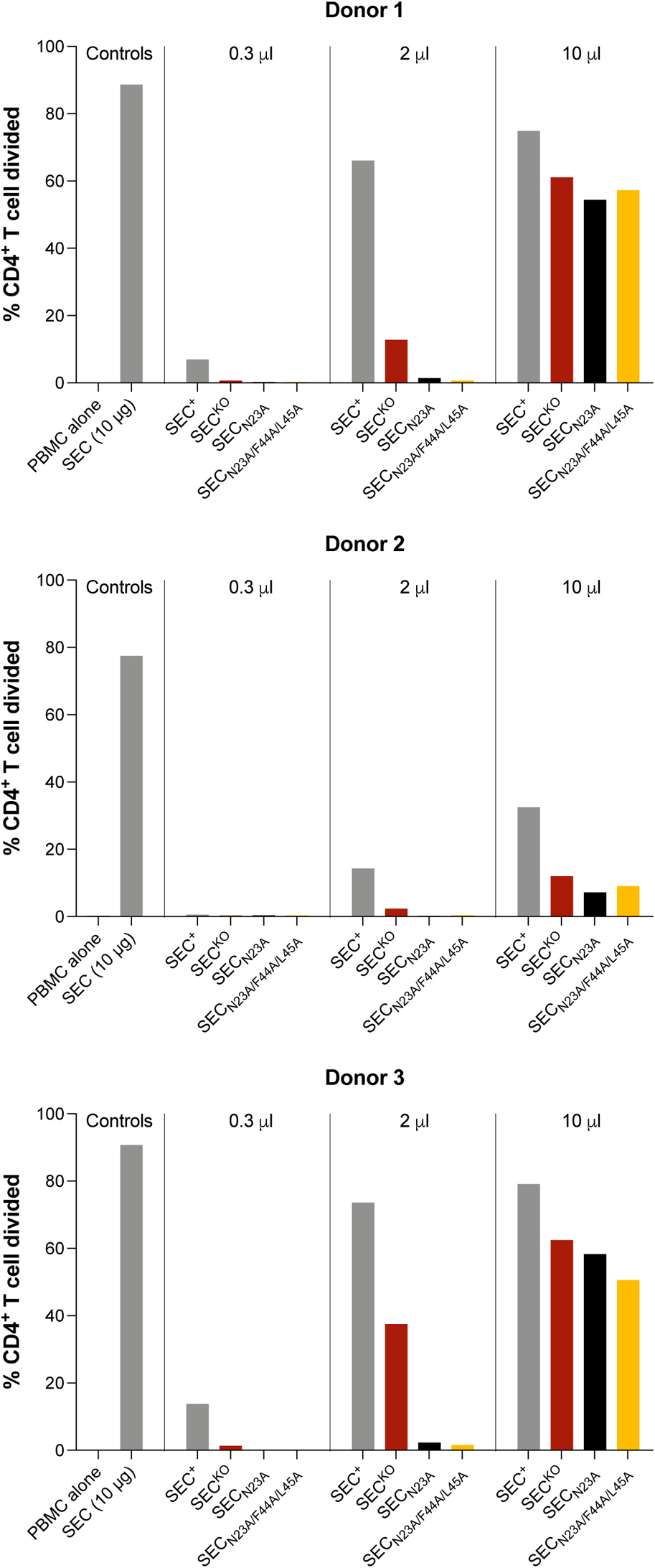
SEC toxoids are deficient in superantigen activity. Percent divided of CD4^+^ T cell from three different human donors. Peripheral blood mononuclear cells (PBMCs) stimulated for 6 days with 0.3, 2, and 10 μl of cell-free supernates from overnight cultures of *S. aureus* SEC^KO^, SEC^+^, SEC_N23A_, and SEC_N23A/F44A/L45A_. PBMCs ± 10 μg of purified SEC were used as controls.

**Figure S3.**
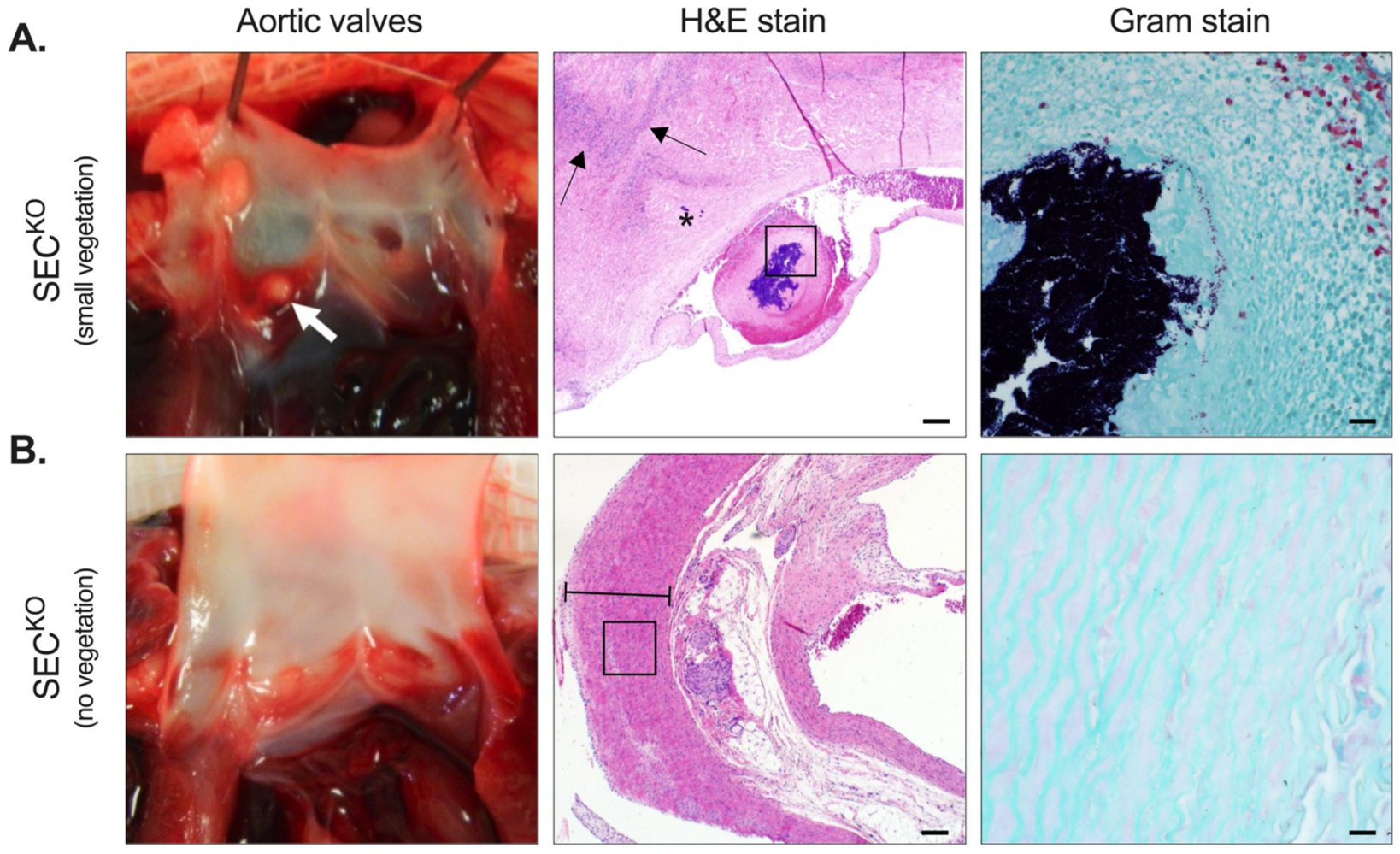
Vegetation formation in rabbits infected with *S. aureus* SEC^KO^. (**A**) Representative image of the aortic valves from *S. aureus* SEC^KO^ infected rabbits that had small proliferative vegetations (white arrow). H&E stained section of a small vegetation (2.6 mm^2^) on the aortic wall with multifocal inflammation throughout the myocardium (arrows) with myocardial infarction adjacent to valvular lesion (star) as well as intralesional bacteria. High magnification Gram stain highlights presence of gram-positive cocci in vegetation. (**B**) Representative image of the aortic valves from *S. aureus* SEC^KO^ infected rabbits that had no proliferative vegetations. H&E stained section of the aortic valve showing no proliferative vegetation, myocardial inflammation, nor visible lesion (perpendicular lines mark the wall of the aorta). High magnification Gram stain image shows absence of bacteria along the myocardium. H&E bar = 200 μm; Gram bar = 20 μm.

**Figure S4.**
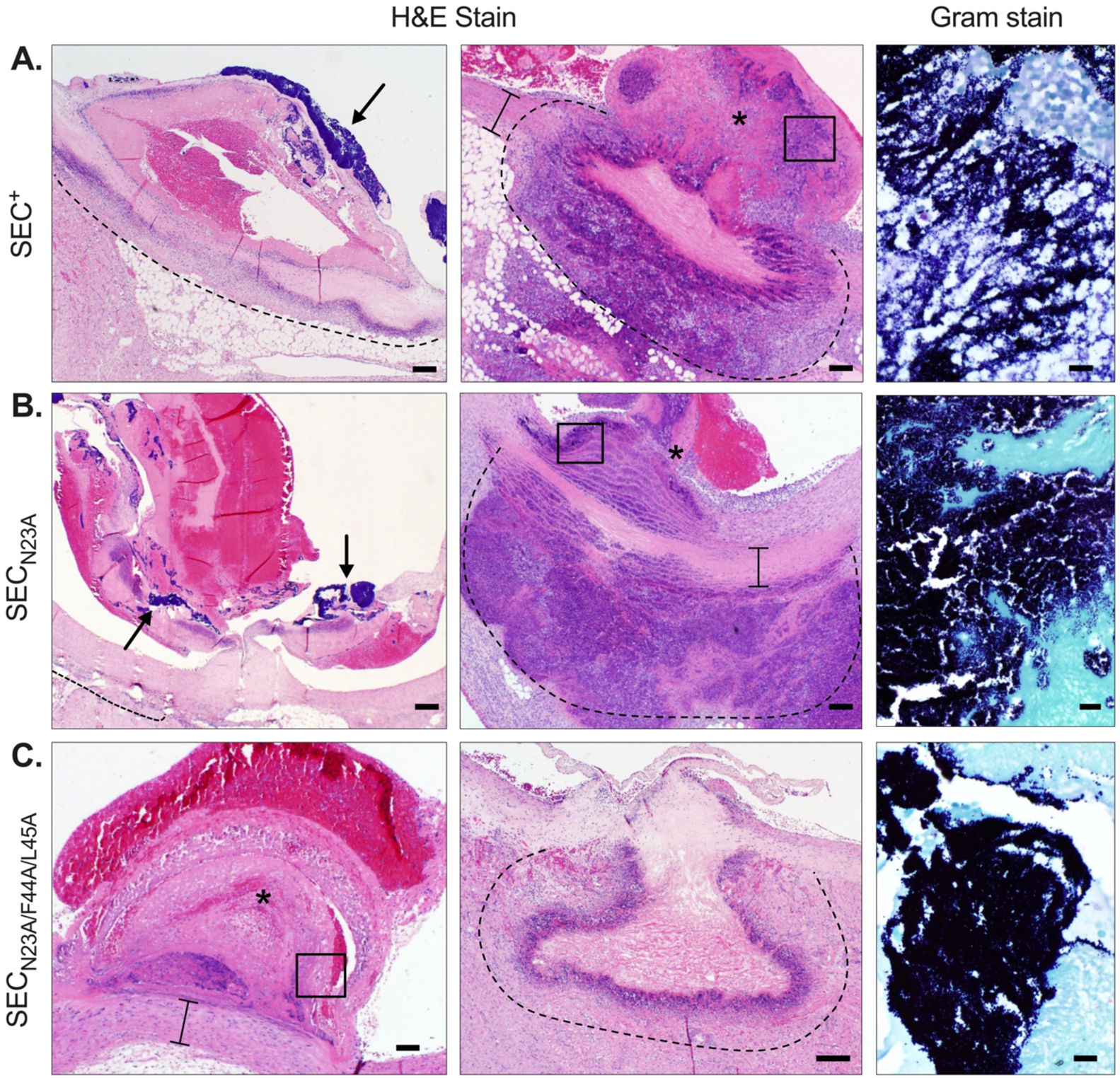
Vegetation formation in rabbits infected with *S. aureus* producing SEC. Images of aortic vegetations from rabbits infected with (**A**) *S. aureus* SEC^+^ (4.42 mm^2^), (**B**) *S. aureus* SEC_N23A_ (10.5 mm^2^), or (**C**) *S. aureus* SEC_N23A/F44A/L45A_ (1.4 mm^2^). *Left panels*: H&E stained sections of aortic valve vegetations showing large bacterial clusters on the aortic endothelium (arrows) and a central core of organized fibrin intermixed with erythrocytes, bacterial colonies and debris. (*) marks the proliferative vegetation on **C**. Bands of heterophils adjacent to the myocardium is shown above the dashed lines (**A-B**) and multifocal zones of heterophilic inflammation along the aortic wall (marked with perpendicular lines) is shown in **C**. *Middle panels*: Images of a proliferative vegetations (star) extending transmurally through the aortic wall (perpendicular lines) with large zones of heterophilic inflammation. (**C**) Image shows vessel wall with necrotic cell debris and fibrin with areas of inflammation within the myocardium shown above the dashed lines. *Right panels*: (**D**) High magnification Gram stain images of a proliferative vegetation from boxed regions in H&E stain showing gram-positive cocci. H&E Bars = 200 μm; Gram bars = 20 μm.

**Figure S5.**
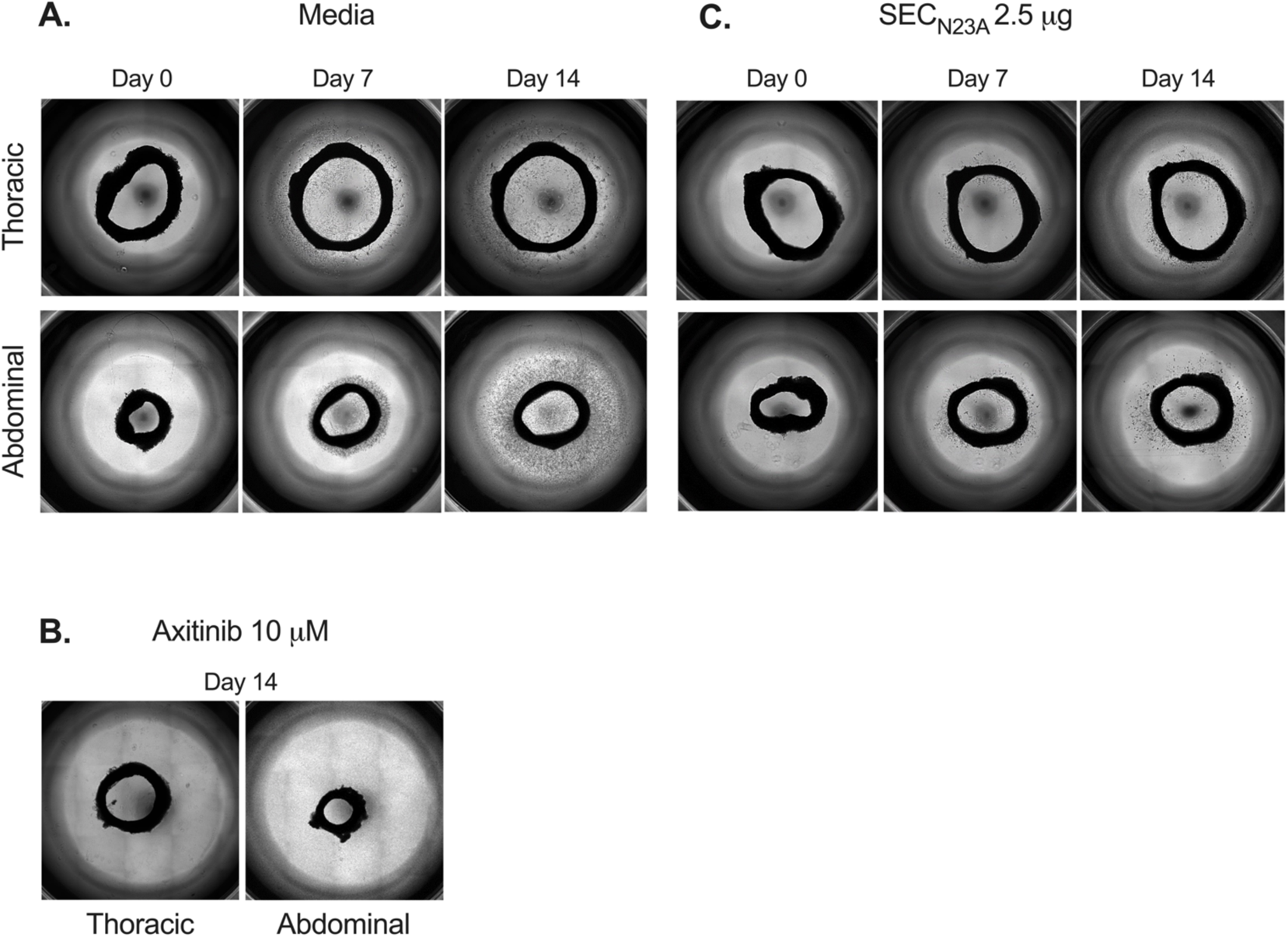
SEC_N23A_ inhibits aortic ring angiogenic sprouting. Thoracic and abdominal aortic ring sections (~1mm wide) derived from 2-3 kg New Zealand white rabbits were embedded in Matrigel basement membrane matrix and cultured in (**A**) media only, (**B**) axitinib (10 μM), or (**C**) SEC_N23A_ (2.5 μg). Sprout formation was captured on day 0, 7, and 14.

**Figure S6.**
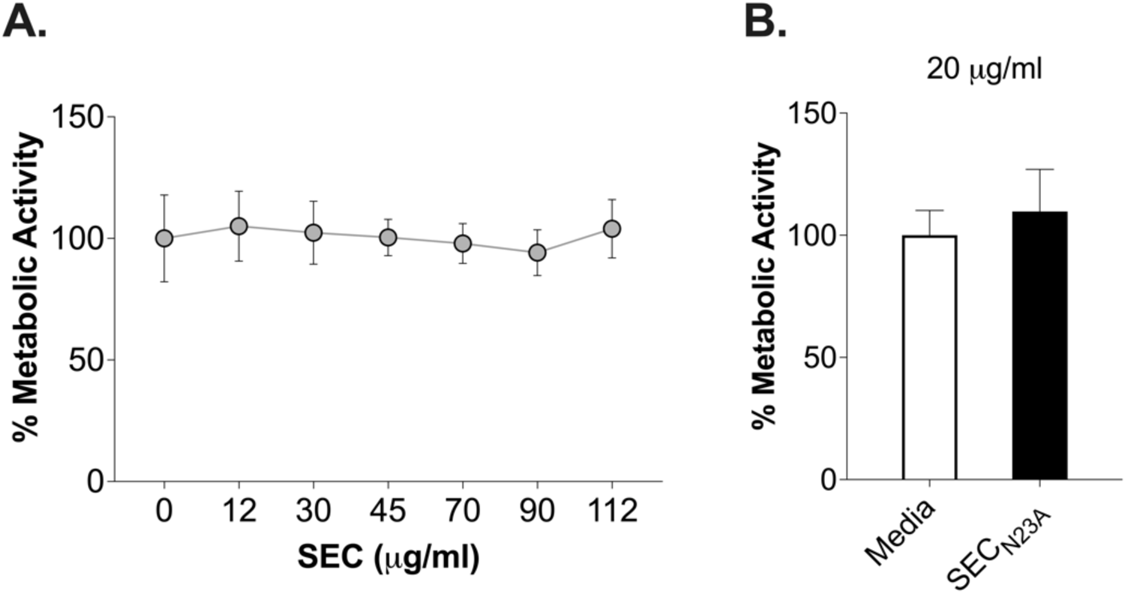
SEC exhibits no cytotoxicity towards immortalized human aortic endothelial cells (iHAECs). Percent metabolic activity of iHAECs grown to near confluency on 1% gelatin-coated plates and treated for 24 h with (**A**) increasing concentrations of SEC (12 – 112 μg mL^−1^) or (**B**) SEC_N23A_ at the experimental dose (20 μg mL^−1^). No statistical significance across groups, one-way ANOVA with the Holm-Šídák’s multiple comparisons test.

## Supplementary Tables

**Table S1.**
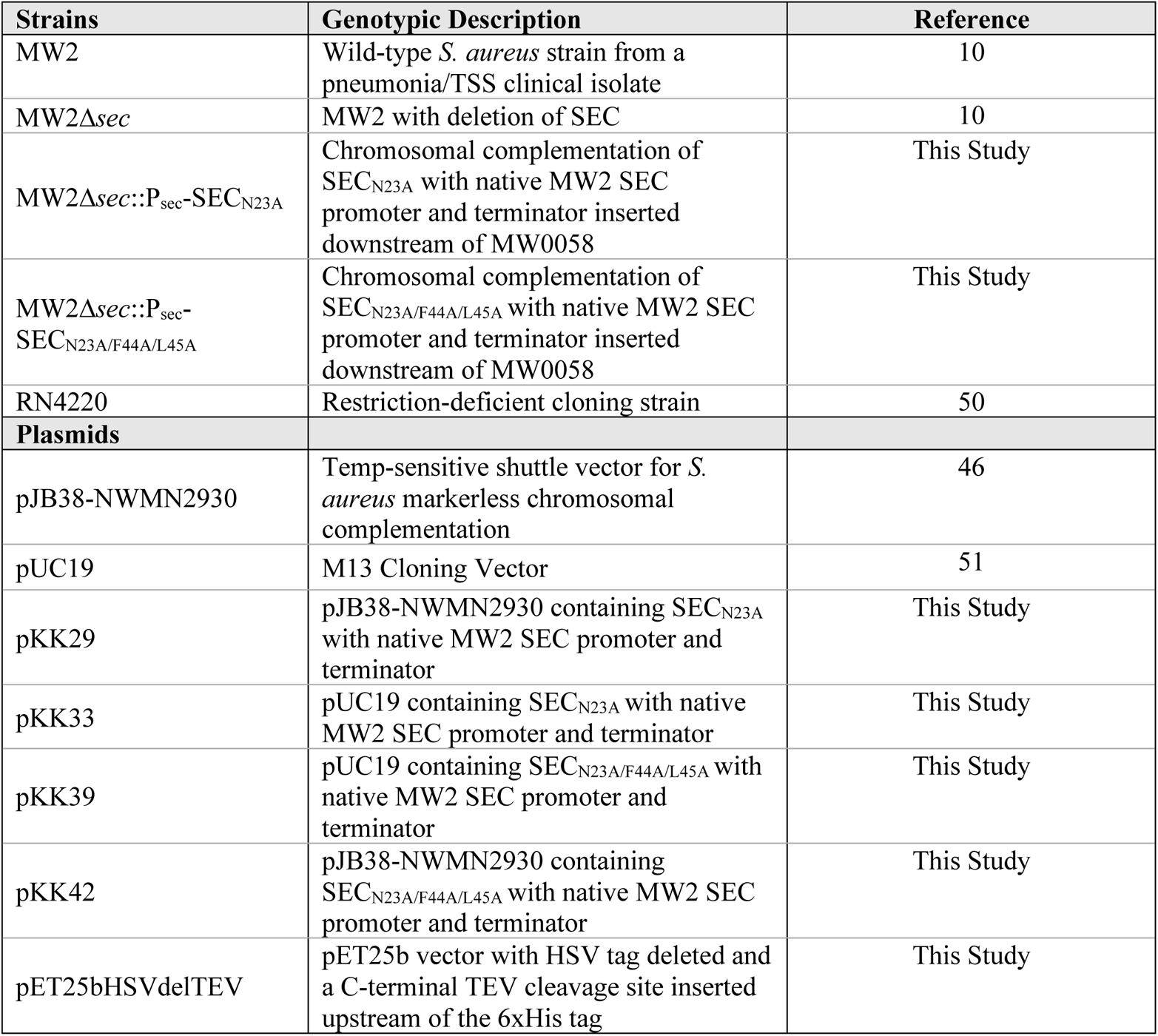
Strains and plasmids.

**Table S2.**
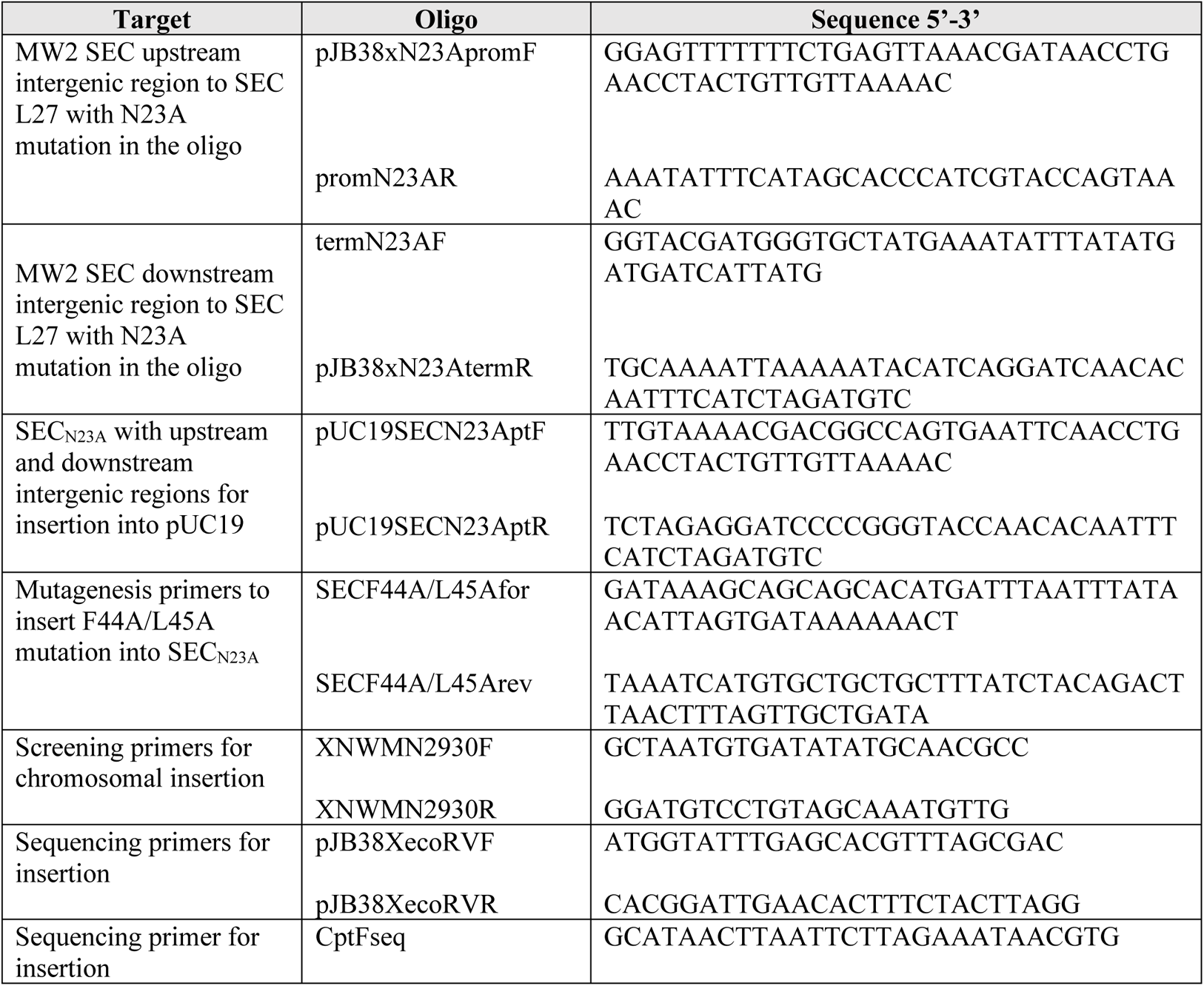
Mutagenesis Primers.

**Table S3.**
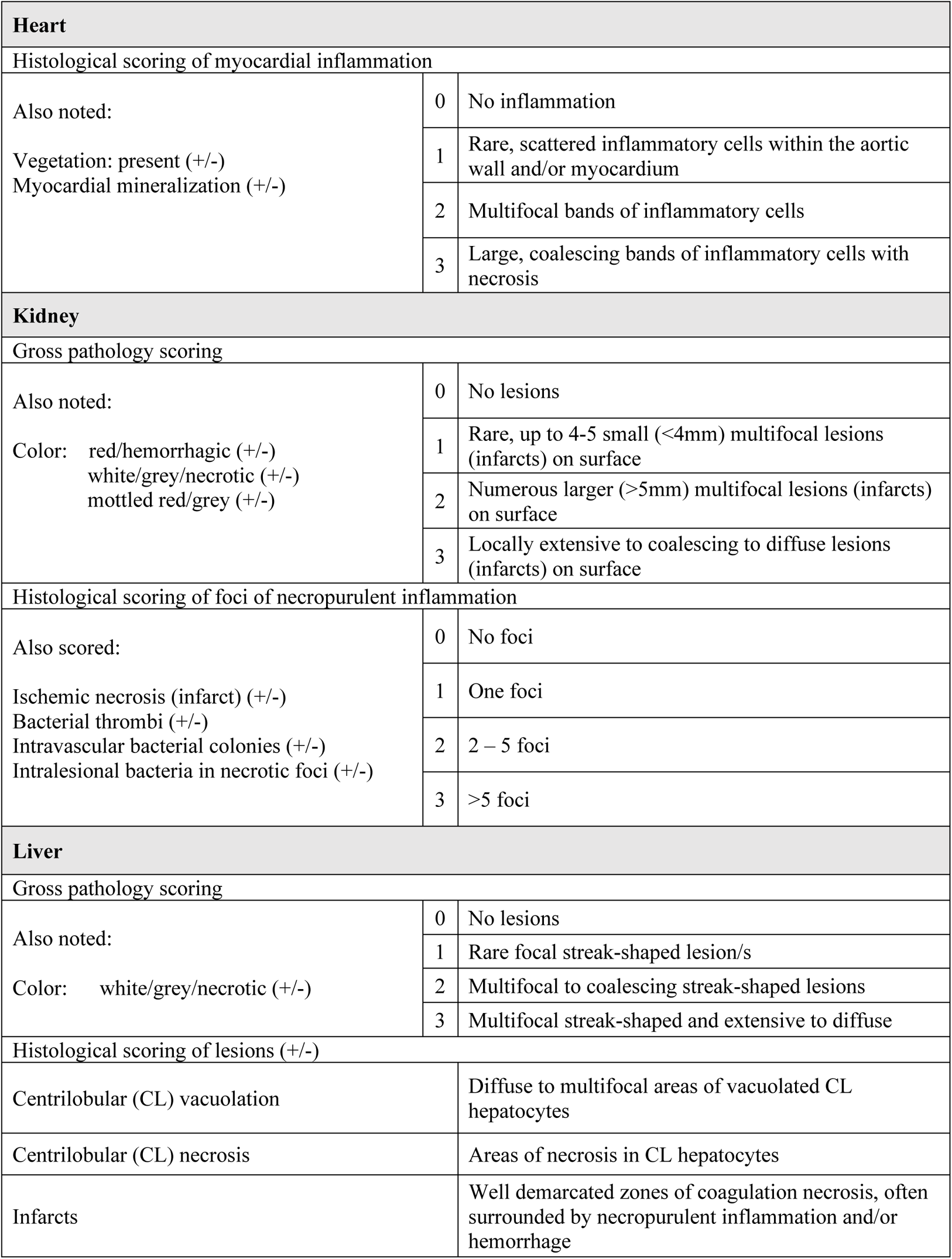
Grading scales and histopathological analyses for the rabbit model of native valve infective endocarditis.

**Table S4.**
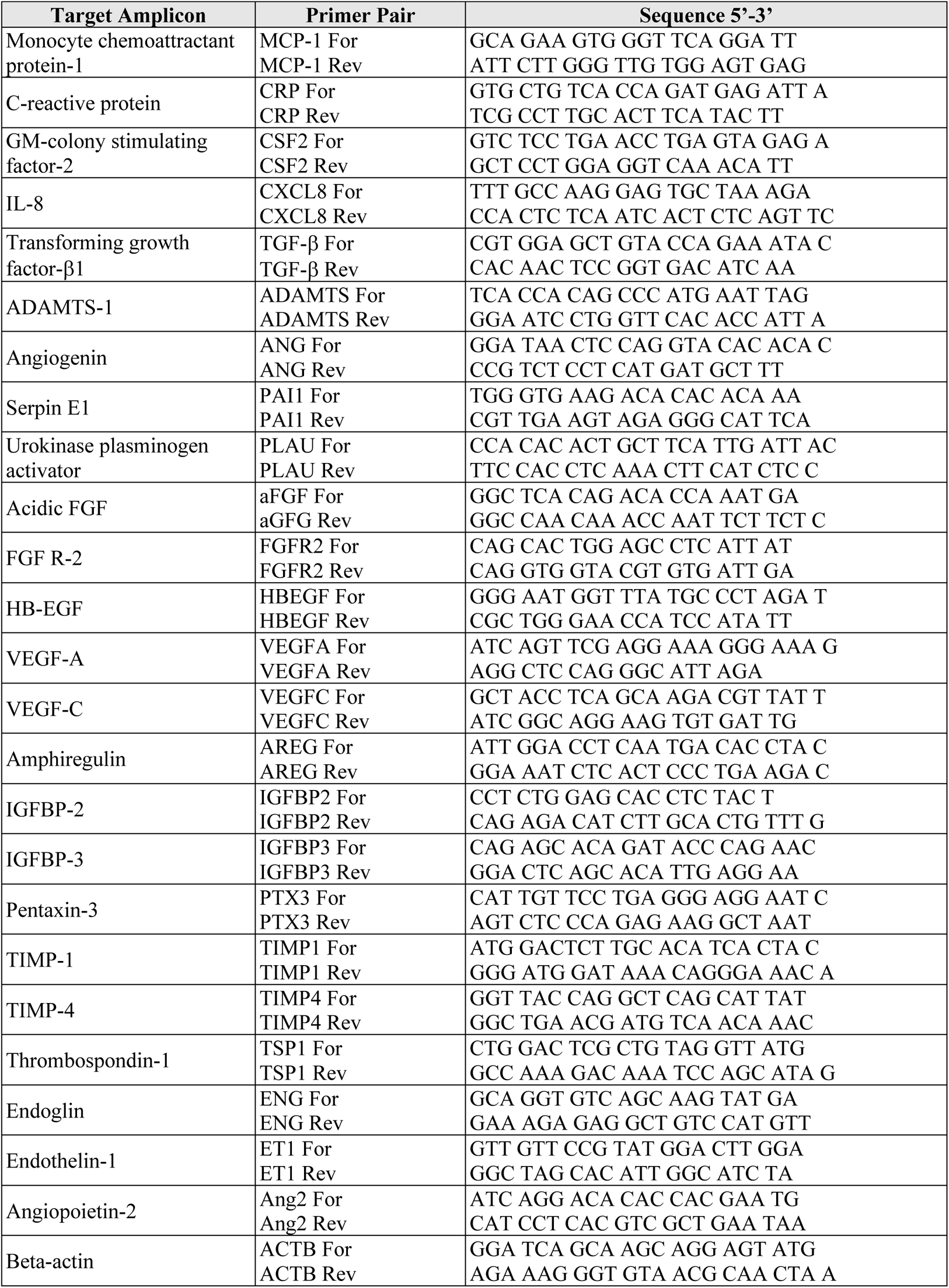
RT-qPCR Primers.

## Notes

### Competing Interest Statement

The authors have declared no competing interest.

### Summary of Updates

We show a novel function of SEC as an anti-angiogenic virulence factor that can exert its effects by inhibiting VEGF-A expression in endothelial cells, explaining SEC's suppression of microvessel formation in the aortic ring model. The overall impact of SEC in IE is now evident as defects in re-endothelialization lead to the progressive loss of the endothelial layer, continual deposition of host factors (e.g. platelets, fibrin, leukocytes), and rapid vegetation growth, while inhibition of sprouting of new capillaries impairs re-vascularization of injured tissues promoting deep tissue toxicities.

## References

1. V. G. Fowler, J. M. Miro, B. Hoen, C. H. Cabell, E. Abrutyn, E. Rubinstein, G. R. Corey, D. Spelman, S. F. Bradley, B. Barsic, P. A. Pappas, K. J. Anstrom, D. Wray, C. Q. Fortes, I. Anguera, E. Athan, P. Jones, J. T. M. Van Der Meer, T. S. J. Elliott, D. P. Levine, A. S. Bayer, *Staphylococcus aureus* endocarditis: A consequence of medical progress, J. Am. Med. Assoc. 293, 3012–3021 (2005).

2. D. H. Bor, S. Woolhandler, R. Nardin, J. Brusch, D. U. Himmelstein, Infective Endocarditis in the U.S., 1998-2009: A Nationwide Study, PLoS One 8, e60033–e60033 (2013).

3. S. Y. C. Tong, J. S. Davis, E. Eichenberger, T. L. Holland, V. G. Fowler, *Staphylococcus aureus* infections: Epidemiology, pathophysiology, clinical manifestations, and management, Clin. Microbiol. Rev. 28, 603–661 (2015).

4. V. G. Fowler, D. T. Durack, Infective endocarditis, Curr. Opin. Cardiol. 9 (1994).

5. G. Thiene, C. Basso, Pathology and pathogenesis of infective endocarditis in native heart valves, Cardiovasc. Pathol. 15, 256–263 (2006).

6. M. L. F. Guerrero, J. J. G. López, A. Goyenechea, J. Fraile, M. De Gó rgolas, Endocarditis caused by *Staphylococcus aureus* a reappraisal of the epidemiologic, clinical, and pathologic manifestations with analysis of factors determining outcome, Medicine (Baltimore). 88, 1–22 (2009).

7. C. Olmos, I. Vilacosta, C. Fernández, J. López, C. Sarriá, C. Ferrera, A. Revilla, J. Silva, D. Vivas, I. González, J. San Román, Contemporary epidemiology and prognosis of septic shock in infective endocarditis, Eur. Heart J. 34, 1999–2006 (2013).

8. E. Klein, D. L. Smith, R. Laxminarayan, Hospitalizations and deaths caused by methicillin-resistant Staphylococcus aureus, United States, 1999-2005, Emerg. Infect. Dis. 13, 1840–1846 (2007).

9. J. J. C. Nienaber, B. K. Sharma Kuinkel, M. Clarke-Pearson, S. Lamlertthon, L. Park, T. H. Rude, S. Barriere, C. W. Woods, V. H. Chu, M. Marín, S. Bukovski, P. Garcia, G. R. Corey, T. Korman, T. Doco-Lecompte, D. R. Murdoch, L. B. Reller, V. G. Fowler, Methicillin-susceptible *Staphylococcus aureus* endocarditis isolates Are associated with clonal complex 30 genotype and a distinct repertoire of enterotoxins and adhesins, J. Infect. Dis. 204, 704–713 (2011).

10. W. Salgado-Pabón, L. Breshears, A. R. Spaulding, J. A. Merriman, C. S. Stach, A. R. Horswill, M. L. Peterson, P. M. Schlievert, E. J. Johnson, Ed. Superantigens are critical for *Staphylococcus aureus* infective endocarditis, sepsis, and acute kidney injury, MBio 4, e00494–13 (2013).

11. C. S. Stach, B. G. Vu, J. A. Merriman, A. Herrera, M. P. Cahill, P. M. Schlievert, W. Salgado-Pabón, Novel Tissue Level Effects of the *Staphylococcus aureus* Enterotoxin Gene Cluster Are Essential for Infective Endocarditis, PLoS One 11, e0154762–e0154762 (2016).

12. A. R. Spaulding, W. Salgado-Pabón, P. L. Kohler, A. R. Horswill, D. Y. M. Leung, P. M. Schlievert, Staphylococcal and streptococcal superantigen exotoxins, Clin. Microbiol. Rev. 26, 422–447 (2013).

13. P. A. Brogan, V. Shah, N. Klein, M. J. Dillon, Vβ-restricted T cell adherence to endothelial cells: A mechanism for superantigen-dependent vascular injury, Arthritis Rheum. 50, 589–597 (2004).

14. S. W. Tuffs, S. M. M. Haeryfar, J. K. McCormick, Manipulation of Innate and Adaptive Immunity by Staphylococcal Superantigens, Pathog. (Basel, Switzerland) 7, 53 (2018).

15. Centers for Disease Control and Prevention, (CDC), Federal Select Agent Program, “Select Agents and Toxins Exclusions: Nontoxic HHS toxins” (Section 73.3 (d)(2); https://www.selectagents.gov/sat/exclusions/nontoxic.htm) [the easiest access to this source is via the URL].

16. A. R. Spaulding, W. Salgado-Pabón, J. A. Merriman, C. S. Stach, Y. Ji, A. N. Gillman, M. L. Peterson, P. M. Schlievert, Vaccination Against *Staphylococcus aureus* Pneumonia, J. Infect. Dis. 209, 1955–1962 (2013).

17. J. Y. Choi, S. Shin, N. Y. Kim, W. S. Son, T. J. Kang, D. H. Song, C. H. Yu, G. H. Hur, S. T. Jeong, Y. K. Shin, A novel staphylococcal enterotoxin B subunit vaccine candidate elicits protective immune response in a mouse model, Toxicon 131, 68–77 (2017).

18. R. G. Ulrich, M. A. Olson, S. Bavari, Development of engineered vaccines effective against structurally related bacterial superantigens, Vaccine 16, 1857–1864 (1998).

19. M. A. Woody, T. Krakauer, B. G. Stiles, Staphylococcal enterotoxin B mutants (N23K and F44S): Biological effects and vaccine potential in a mouse model, Vaccine 15, 133–139 (1997).

20. T. S. Jardetzky, J. H. Brown, J. C. Gorga, L. J. Stern, R. G. Urban, Y. I. Chi, C. Stauffacher, J. L. Strominger, D. C. Wiley, Three-dimensional structure of a human class II histocompatibility molecule complexed with superantigen, Nature 368, 711–718 (1994).

21. M. A. Woody, T. Krakauer, R. G. Ulrich, B. G. Stiles, Differential Immune Responses to Staphylococcal Enterotoxin B Mutations in a Hydrophobic Loop Dominating the Interface with Major Histocompatibility Complex Class II Receptors, J. Infect. Dis. 177, 1013–1022 (1998).

22. P. A. Szabo, A. Goswami, D. M. Mazzuca, K. Kim, D. B. O’Gorman, D. A. Hess, I. D. Welch, H. A. Young, B. Singh, J. K. McCormick, S. M. M. Haeryfar, Rapid and Rigorous IL-17A Production by a Distinct Subpopulation of Effector Memory T Lymphocytes Constitutes a Novel Mechanism of Toxic Shock Syndrome Immunopathology, J. Immunol. 198, 2805–2818 (2017).

23. J. G. Zein, G. L. Lee, M. Tawk, M. Dabaja, G. T. Kinasewitz, Prognostic Significance of Elevated Serum Lactate Dehydrogenase (LDH) in Patients with Severe Sepsis, Chest 126, 873S (2004).

24. K. Hayashida, Y. Chen, A. H. Bartlett, W. P. Pyong, Syndecan-1 is an *in vivo* suppressor of gram-positive toxic shock, J. Biol. Chem. 283, 19895–19903 (2008).

25. R. Sada, S. Fukuda, H. Ishimaru, Toxic shock syndrome due to community-acquired methicillin-resistant *Staphylococcus aureus* infection: Two case reports and a literature review in Japan, IDCases 8, 77–80 (2017).

26. E. G. Giannini, R. Testa, V. Savarino, Liver enzyme alteration: a guide for clinicians, CMAJ 172, 367–379 (2005).

27. V. Fuhrmann, B. Jäger, A. Zubkova, A. Drolz, Hypoxic hepatitis – epidemiology, pathophysiology and clinical management, Wien. Klin. Wochenschr. 122, 129–139 (2010).

28. C. N. Gentry, J. R. McDonald, Acute infective endocarditis, Infect. Dis. Crit. Care 23, 271–283 (2007).

29. T. R. Wojda, K. Cornejo, A. Lin, A. Cipriano, S. Nanda, J. D. Amortegui, B. T. Wojda, S. P. Stawicki, “Septic Embolism: A Potentially Devastating Complication of Infective Endocarditis” in Contemporary Challenges in Endocarditis, M.S. Firstenberg, Eds. (IntechOpen, 2016), chap. 8, DOI: 10.5772/64931.

30. T. H. Adair and J.-P. Montani, “Colloquium series on integrated systems physiology: from molecule to function to disease,” in Angiogenesis, D. N. Granger and J. P. Granger, Eds (Morgan & Claypool, 2010), Vol. 2, No. 1, Pages 1–84.

31. F. Moccia, S. Negri, M. Shekha, P. Faris, G. Guerra, Endothelial Ca^2+^ signaling, angiogenesis and vasculogenesis: Just what it takes to make a blood vessel, Int. J. Mol. Sci. 20 (2019).

32. R. J. Kelly, O. Rixe, Axitinib-a selective inhibitor of the vascular endothelial growth factor (VEGF) receptor, Target. Oncol. 4, 297–305 (2009).

33. A. Hoeben, B. Landuyt, M. S. Highley, H. Wildiers, A. T. Van Oosterom, E. A. De Bruijn, Vascular endothelial growth factor and angiogenesis, Pharmacol. Rev. 56, 549–580 (2004).

34. P. Carmeliet, VEGF as a key mediator of angiogenesis in cancer, Oncology 69, 4–10 (2005).

35. P. M. Tran, S. S. Tang, W. Salgado-Pabón, *Staphylococcus aureus* β-Toxin Exerts Anti-angiogenic Effects by Inhibiting Re-endothelialization and Neovessel Formation*Front*. Microbiol. 13 (2022).

36. Y. Li, H. Zhu, X. Wei, H. Li, Z. Yu, H. M. Zhang, W. C. Liu, LPS induces HUVEC angiogenesis *in vitro* through miR-146a-mediated TGF-β1 inhibition, Am. J. Transl. Res. 9, 591–600 (2017).

37. K. Werdan, S. Dietz, B. Löffler, S. Niemann, H. Bushnaq, R. E. Silber, G. Peters, U. Müller-Werdan, Mechanisms of infective endocarditis: Pathogen-host interaction and risk states, Nat. Rev. Cardiol. 11, 35–50 (2014).

38. K. Kulhankova, K. J. Kinney, J. M. Stach, F. A. Gourronc, I. M. Grumbach, A. J. Klingelhutz, W. Salgado-Pabón, The Superantigen Toxic Shock Syndrome Toxin 1 Alters Human Aortic Endothelial Cell Function, Infect. Immun. 86, e00848–17 (2018).

39. A. J. Brosnahan, M. M. Schaefers, W. H. Amundson, M. J. Mantz, C. A. Squier, M. L. Peterson, P. M. Schlievert, Novel toxic shock syndrome toxin-1 amino acids required for biological activity, Biochemistry 47, 12995–13003 (2008).

40. S. W. Tuffs, D. B. A. James, J. Bestebroer, A. C. Richards, M. I. Goncheva, M. O’Shea, B. A. Wee, K. S. Seo, P. M. Schlievert, A. Lengeling, J. A. van Strijp, V. J. Torres, J. R. Fitzgerald, The *Staphylococcus aureus* superantigen SElX is a bifunctional toxin that inhibits neutrophil function, PLoS Pathog. 13 (2017).

41. D. Grumann, S. S. Scharf, S. Holtfreter, C. Kohler, L. Steil, S. Engelmann, M. Hecker, U. Völker, B. M. Bröker, Immune Cell Activation by Enterotoxin Gene Cluster (egc)-Encoded and Non-egc Superantigens from *Staphylococcus aureus*, J. Immunol. 181, 5054–5061 (2008).

42. D. S. Terman, A. Serier, O. Dauwalder, C. Badiou, A. Dutour, D. Thomas, V. Brun, J. Bienvenu, J. Etienne, F. Vandenesch, G. Lina, Staphylococcal entertotoxins of the enterotoxin gene cluster (egcSEs) induce nitric oxide- and cytokine dependent tumor cell apoptosis in a broad panel of human tumor cells, Front. Cell. Infect. Microbiol. 4, 38 (2013).

43. D. A. Chistiakov, A. N. Orekhov, Y. V. Bobryshev, Endothelial barrier and its abnormalities in cardiovascular disease, Front. Physiol. 6, 365 (2015).

44. H. Gerhardt, C. Betsholtz, Endothelial-pericyte interactions in angiogenesis, Cell Tissue Res. 314, 15–23 (2003).

45. D. Ribatti, E. Crivellato, Immune cells and angiogenesis, J. Cell. Mol. Med. 13, 2822–2833 (2009).

46. V. Baeriswyl, G. Christofori, The angiogenic switch in carcinogenesis, Semin. Cancer Biol. 19, 329–337 (2009).

47. H. Takahashi, M. Shibuya, The vascular endothelial growth factor (VEGF)/VEGF receptor system and its role under physiological and pathological conditions, Clin. Sci. 109, 227–241 (2005).

48. J. A. Nagy, A. M. Dvorak, H. F. Dvorak, VEGF-A and the induction of pathological angiogenesis, Annu. Rev. Pathol. 2, 251–275 (2007).

49. J. E. Nör, J. Christensen, D. J. Mooney, P. J. Polverini, Vascular endothelial growth factor (VEGF)-mediated angiogenesis is associated with enhanced endothelial cell survival and induction of Bcl-2 expression, Am. J. Pathol. 154, 375–384 (1999).

50. H. P. Gerber, A. McMurtrey, J. Kowalski, M. Yan, B. A. Keyt, V. Dixit, N. Ferrara, Vascular endothelial growth factor regulates endothelial cell survival through the phosphatidylinositol 3’-kinase/Akt signal transduction pathway: Requirement for Flk-1/KDR activation, J. Biol. Chem. 273, 30336–30343 (1998).

51. R. Hutter, F. E. Carrick, C. Valdiviezo, C. Wolinsky, J. S. Rudge, S. J. Wiegand, V. Fuster, J. J. Badimon, B. V. Sauter, Vascular endothelial growth factor regulates reendothelialization and neointima formation in a mouse model of arterial injury, Circulation 110, 2430–2435 (2004).

52. J. Banbury, M. Siemionow, S. Porvasnik, S. Petras, E. Browne, Improved perfusion after subcritical ischemia in muscle flaps treated with vascular endothelial growth factor, Plast. Reconstr. Surg. 106, 1541–1546 (2000).

53. J. J. Lopez, R. J. Laham, A. Stamler, J. D. Pearlman, S. Bunting, A. Kaplan, J. P. Carrozza, F. W. Sellke, M. Simons, VEGF administration in chronic myocardial ischemia in pigs, Cardiovasc. Res. 40, 272–281 (1998).

54. G. E. Y. Cao, S. Bhattacharya, S. Dutta, E. Wang, D. Mukhopadhyay, Endogenous vascular endothelial growth factor-A (VEGF-A) maintains endothelial cell homeostasis by regulating VEGF receptor-2 transcription, J. Biol. Chem. 287, 3029–3041 (2012).

55. B. K. McColl, S. A. Stacker, M. G. Achen, Molecular regulation of the VEGF family - Inducers of angiogenesis and lymphangiogenesis, Apmis 112, 463–480 (2004).

56. S. Lee, T. T. Chen, C. L. Barber, M. C. Jordan, J. Murdock, S. Desai, N. Ferrara, A. Nagy, K. P. Roos, M. L. Iruela-Arispe, Autocrine VEGF Signaling Is Required for Vascular Homeostasis, Cell 130, 691–703 (2007).

57. J. Siedlecki, C. Wertheimer, A. Wolf, R. Liegl, C. Priglinger, S. Priglinger, K. Eibl-Lindner, Combined VEGF and PDGF inhibition for neovascular AMD: anti-angiogenic properties of axitinib on human endothelial cells and pericytes *in vitro*, Graefe’s Arch. Clin. Exp. Ophthalmol. 255, 963–972 (2017).

58. N. Yamamoto, T. Oyaizu, M. Enomoto, M. Horie, M. Yuasa, A. Okawa, K. Yagishita, VEGF and bFGF induction by nitric oxide is associated with hyperbaric oxygen-induced angiogenesis and muscle regeneration, Sci. Rep. 10, 2744 (2020).

59. R. Hlushchuk, M. Ehrbar, P. Reichmuth, N. Heinimann, B. Styp-Rekowska, R. Escher, O. Baum, P. Lienemann, A. Makanya, E. Keshet, V. Djonov, Decrease in VEGF expression induces intussusceptive vascular pruning, Arterioscler. Thromb. Vasc. Biol. 31, 2836–2844 (2011).

60. D. R. Murdoch, R. G. Corey, B. Hoen, M. Miró, V. G. Fowler, A. S. Bayer, A. W. Karchmer, L. Olaison, P. A. Pappas, P. Moreillon, S. T. Chambers, V. H. Chu, V. Falcó, D. J. Holland, P. Jones, J. L. Klein, N. J. Raymond, K. M. Read, M. F. Tripodi, R. Utili, A. Wang, C. W. Woods, C. H. Cabell, Clinical presentation, etiology, and outcome of infective endocarditis in the 21st century The international collaboration on Endocarditis-prospective cohort study, Arch. Intern. Med. 169, 463–473 (2009).

61. J. W. Shupp, M. Jett, C. H. Pontzer, Identification of a transcytosis epitope on staphylococcal enterotoxins, Infect. Immun. 70, 2178–2186 (2002).

62. B. A. Fields, E. L. Malchiodi, H. Li, X. Ysern, C. V Stauffacher, P. M. Schlievert, K. Karjalainen, R. A. Mariuzza, Crystal structure of a T-cell receptor β-chain complexed with a superantigen, Nature 384, 188–192 (1996).

63. L. Leder, A. Llera, P. M. Lavoie, M. I. Lebedeva, H. Li, R. P. Sékaly, G. A. Bohach, P. J. Gahr, P. M. Schlievert, K. Karjalainen, R. A. Mariuzza, A mutational analysis of the binding of staphylococcal enterotoxins B and C3 to the T cell receptor beta chain and major histocompatibility complex class II, J. Exp. Med. 187, 823–833 (1998).

64. R. Ulrich, S. Bavari, M. Olson, Staphylococcal enterotoxins A and B share a common structural motif for binding class II major histocompatibility complex molecules, Nat. Struct. Biol. 2, 554–560 (1995).

65. J. W. Kappler, A. Herman, J. Clements, P. Marrack, Mutations defining functional regions of the superantigen staphylococcal enterotoxin B, J. Exp. Med. 175, 387–396 (1992).

66. N. W. M. de Jong, T. van der Horst, J. A. G. van Strijp, R. Nijland, Fluorescent reporters for markerless genomic integration in *Staphylococcus aureus*, Sci. Rep. 7, 43889 (2017).

67. D. G. Gibson, L. Young, R.-Y. Chuang, J. C. Venter, C. A. Hutchison III, H. O. Smith, Enzymatic assembly of DNA molecules up to several hundred kilobases, Nat. Methods 6, 343 (2009).

68. M. R. Grosser, A. R. Richardson, in Methods in Molecular Biology, J. L. Bose, Ed. (Springer New York, New York, NY, 2016), vol. 1373, pp. 51–57.

69. J. A. Merriman, P. M. Schlievert, in Methods in Molecular Biology, A. J. Brosnahan, Ed. (Springer New York, New York, NY, 2016), vol. 1396, pp. 19–33.

70. K. N. Gibson-Corley, A. K. Olivier, D. K. Meyerholz, Principles for Valid Histopathologic Scoring in Research, Vet. Pathol. 50, 1007–1015 (2013).

71. W.-H. Zhu, R. F. Nicosia, The thin prep rat aortic ring assay: A modified method for the characterization of angiogenesis in whole mounts, Angiogenesis 5, 81–86 (2002).

72. J. Stiffey-Wilusz, J. A. Boice, J. Ronan, A. M. Fletcher, M. S. Anderson, An ex vivo angiogenesis assay utilizing commercial porcine carotid artery: Modification of the rat aortic ring assay, Angiogenesis 4, 3–9 (2001).

73. J. M. King, K. Kulhankova, C. S. Stach, B. G. Vu, W. Salgado-Pabón, Phenotypes and Virulence among Staphylococcus aureus USA100, USA200, USA300, USA400, and USA600 Clonal Lineages, mSphere 1, e00071–16 (2016).

