## supplementary materials for the preprint site for "SEC is an anti-angiogenic virulence factor that promotes *Staphylococcus aureus* Infective Endocarditis Independent of Superantigen Activity"

### Supplementary Materials and Methods

#### *S. aureus* growth curves

Overnight cultures were diluted to an $OD_{600}$ 1.0 in TH. 10 µl of diluted culture was added to 190 µl of TH in a 96-well round, clear bottom plate in technical triplicate (Corning). Plates were sealed using Breathe-Easy sealing membrane (Diversified Biotech). Growth was measured using a TECAN M200 plate reader overnight with the following settings: 37°C, 30 min kinetic interval, orbital shaking: amplitude 3 mm for 10 s before reads, 1 read/well at 600 nM, 25 flashes, 9 nM bandwidth, 100ms settle time, and 27 min wait time between reads with orbital shaking: amplitude 3mm for 26.5 min. Data is represented by three independent experiments.

#### Erythrocyte lysis assays for hemolysin production

Erythrocyte Lysis assays were performed as previously described *(73)*. Overnight cultures were diluted to an $OD_{600}$ 1.0 in PBS and 5 µl spotted onto TSA II agar plates with either 5% rabbit or sheep blood (Becton Dickinson). Plates were incubated for 24 h at 37°C with 5% $CO_2$. Zones of hemolysis were measured and quantified using ImageJ. Data is represented by three independent experiments done with technical duplicates.

#### MTS assay

An MTS assay was used to determine cell viability. Cells were seeded at 7,000 cells/well into gelatin-coated 96-well plates and grown overnight to near confluency. Media was removed and replaced with 100 µL of media containing increasing concentrations of SEC or $SEC_{N23A}$ at 20 µg $mL^{-1}$ followed by overnight incubation. 20 µL of CellTiter 96® AQueous One Solution was added to each well followed by a 1 h incubation at 37°C, 5% $CO_2$. A plate reader was used to read absorbance at 490 nm. Three biological replicates were conducted with each containing three technical replicates. The data were normalized so that untreated cells were considered 100% activity by dividing the absorbance of treated cells by the absorbance of untreated cells.

### Supplementary figures

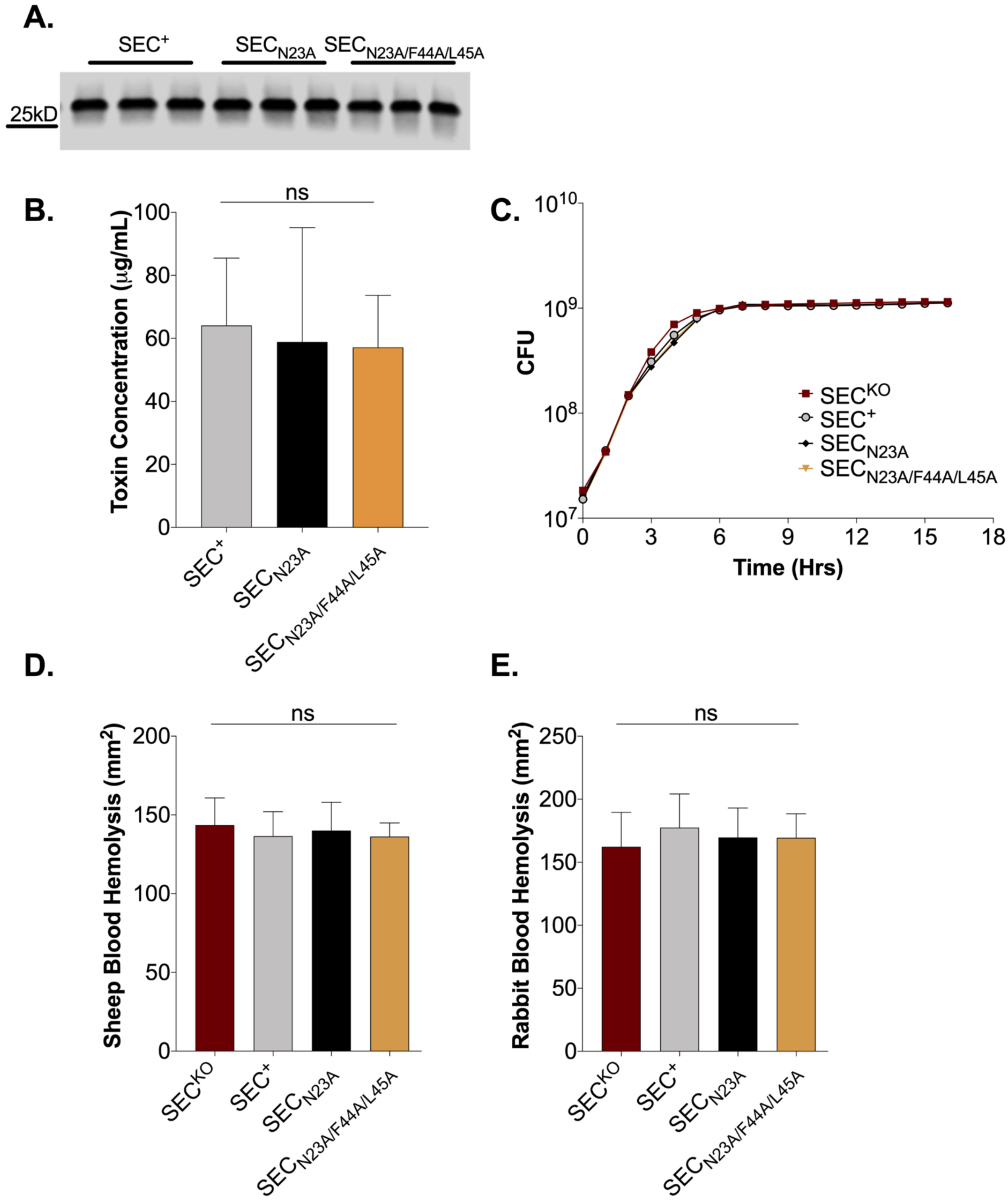

**Figure S1. Virulence factor production and growth of strains producing SEC binding site inactivated toxoids.** (**A**) Representative western blot images of SEC production in wild-type and SEC complement strains grown overnight in Todd Hewitt (TH) medium shown in technical triplicate. (**B**) Quantitative western blot analysis of SEC production. (**C**) Growth curve of *S. aureus* $SEC^{KO}$, $SEC^{+}$, $SEC_{N23A}$, $SEC_{N23A/F44A/L45A}$ grown overnight in TH. (**D**) Relative levels of hemolysin production as measured in a sheep erythrocyte lysis assay. (**E**) Relative levels of α-hemolysin production as measured in a rabbit erythrocyte lysis assay. (**D, E**) Overnight cultures of *S. aureus* were washed and spotted onto blood agar plates. Zones of hemolysis were measured after overnight growth. Data are represented as mean (± SD). Statistical significance was determined by one-way ANOVA and nonparametric Kruskal-Wallis test. $p$ values $\leq 0.05$ are considered statistically significant.

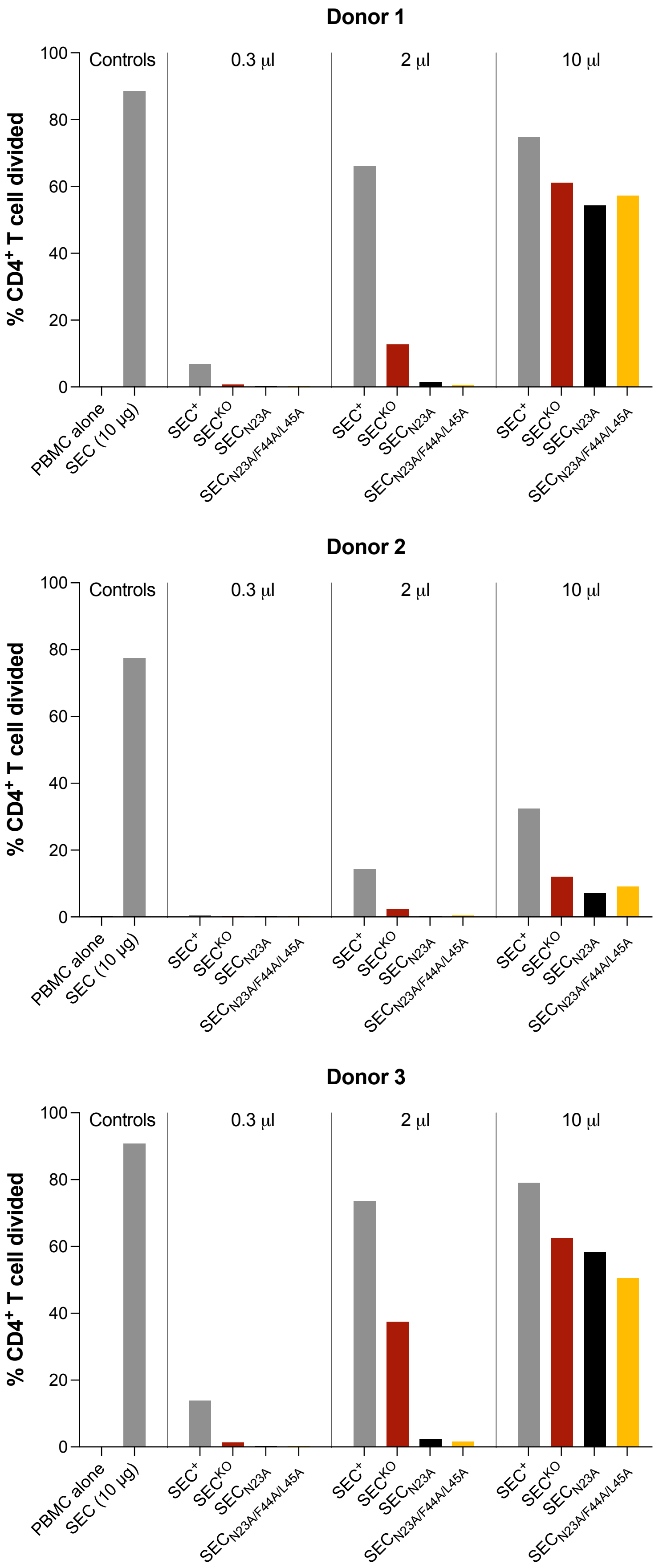
Donor 1
Donor 2
Donor 3
% CD4+ T cell divided
Controls
0.3 µl
2 µl
10 µl
0
20
40
60
80
100
PBMC alone
SEC (10 µg)
SEC+
SECKO
SECN23A
SECN23A/F44A/L45A

**Figure S2. SEC toxoids are deficient in superantigen activity.** Percent divided of $CD4^+$ T cell from three different human donors. Peripheral blood mononuclear cells (PBMCs) stimulated for 6 days with 0.3, 2, and 10 µl of cell-free supernates from overnight cultures of *S. aureus* $SEC^{KO}$, $SEC^{+}$, $SEC_{N23A}$, and $SEC_{N23A/F44A/L45A}$. PBMCs ± 10 µg of purified SEC were used as controls.

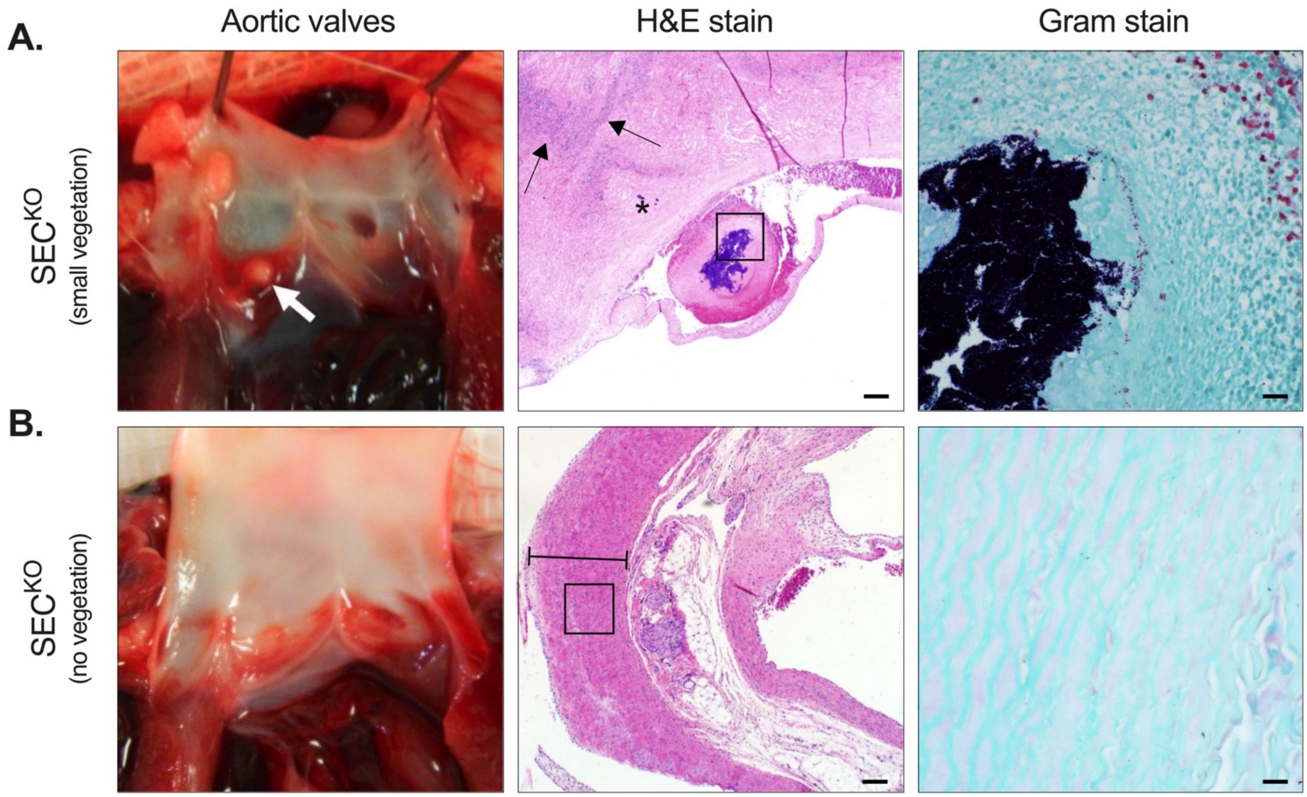


**Figure S3. Vegetation formation in rabbits infected with *S. aureus* SEC$^{KO}$.** (**A**) Representative image of the aortic valves from *S. aureus* SEC$^{KO}$ infected rabbits that had small proliferative vegetations (white arrow). H&E stained section of a small vegetation (2.6 mm$^2$) on the aortic wall with multifocal inflammation throughout the myocardium (arrows) with myocardial infarction adjacent to valvular lesion (star) as well as intralesional bacteria. High magnification Gram stain highlights presence of gram-positive cocci in vegetation. (**B**) Representative image of the aortic valves from *S. aureus* SEC$^{KO}$ infected rabbits that had no proliferative vegetations. H&E stained section of the aortic valve showing no proliferative vegetation, myocardial inflammation, nor visible lesion (perpendicular lines mark the wall of the aorta). High magnification Gram stain image shows absence of bacteria along the myocardium. H&E bar = 200 µm; Gram bar = 20 µm.

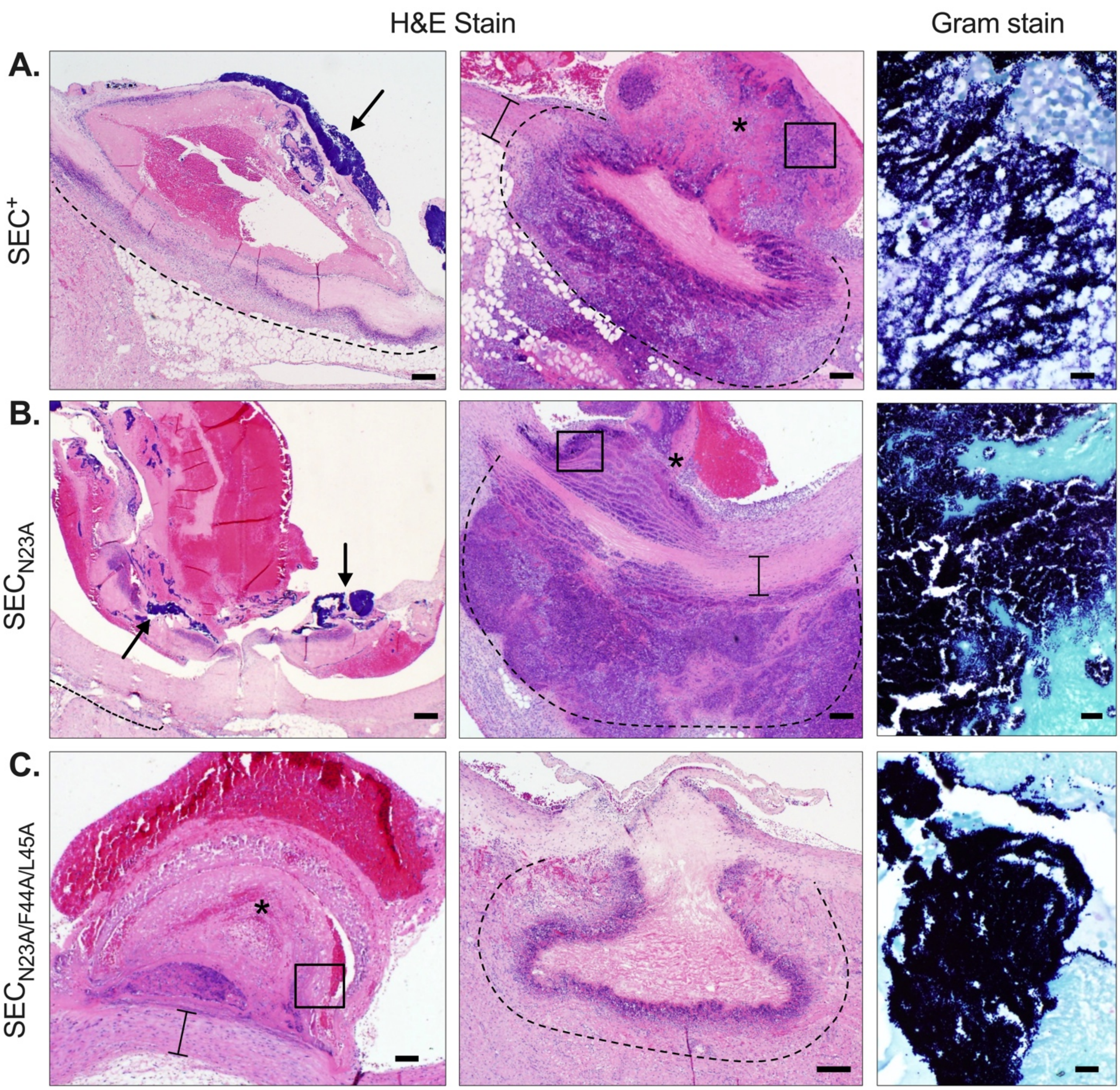


**Figure S4. Vegetation formation in rabbits infected with *S. aureus* producing SEC.** Images of aortic vegetations from rabbits infected with (**A**) *S. aureus* SEC$^+$ (4.42 mm$^2$), (**B**) *S. aureus* SEC$_{N23A}$ (10.5 mm$^2$), or (**C**) *S. aureus* SEC$_{N23A/F44A/L45A}$ (1.4 mm$^2$). *Left panels*: H&E stained sections of aortic valve vegetations showing large bacterial clusters on the aortic endothelium (arrows) and a central core of organized fibrin intermixed with erythrocytes, bacterial colonies and debris. (*) marks the proliferative vegetation on **C**. Bands of heterophils adjacent to the myocardium is shown above the dashed lines (**A-B**) and multifocal zones of heterophilic inflammation along the aortic wall (marked with perpendicular lines) is shown in **C**. *Middle panels*: Images of a proliferative vegetations (star) extending transmurally through the aortic wall (perpendicular lines) with large zones of heterophilic inflammation. (**C**) Image shows vessel wall with necrotic cell debris and fibrin with areas of inflammation within the myocardium shown above the dashed lines. *Right panels*: (**D**) High magnification Gram stain images of a proliferative vegetation from boxed regions in H&E stain showing gram-positive cocci. H&E Bars = 200 μm; Gram bars = 20 μm.

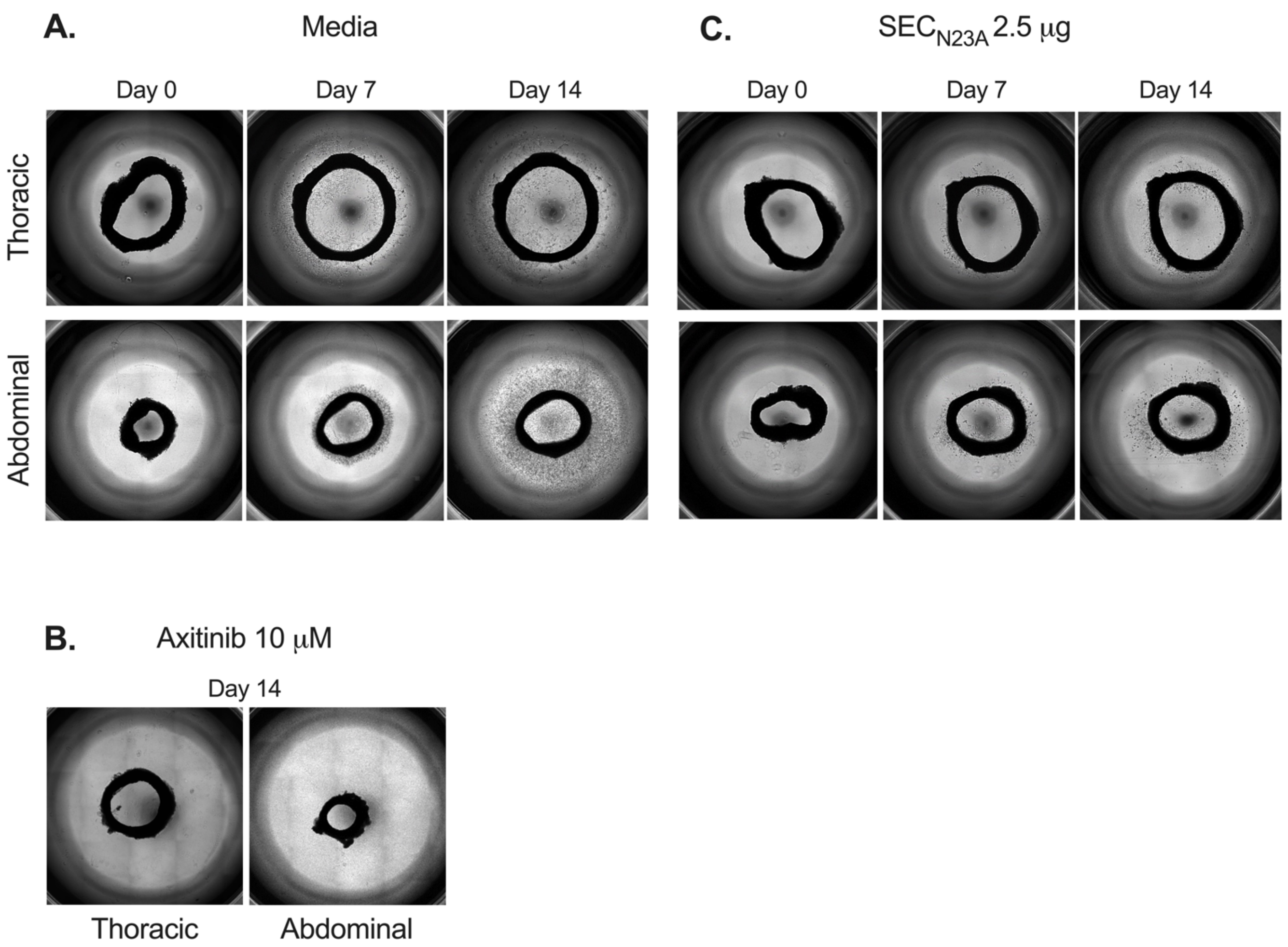


**Figure S5. $SEC_{N23A}$ inhibits aortic ring angiogenic sprouting.** Thoracic and abdominal aortic ring sections (~1mm wide) derived from 2-3 kg New Zealand white rabbits were embedded in Matrigel basement membrane matrix and cultured in (**A**) media only, (**B**) axitinib (10 μM), or (**C**) $SEC_{N23A}$ (2.5 μg). Sprout formation was captured on day 0, 7, and 14.

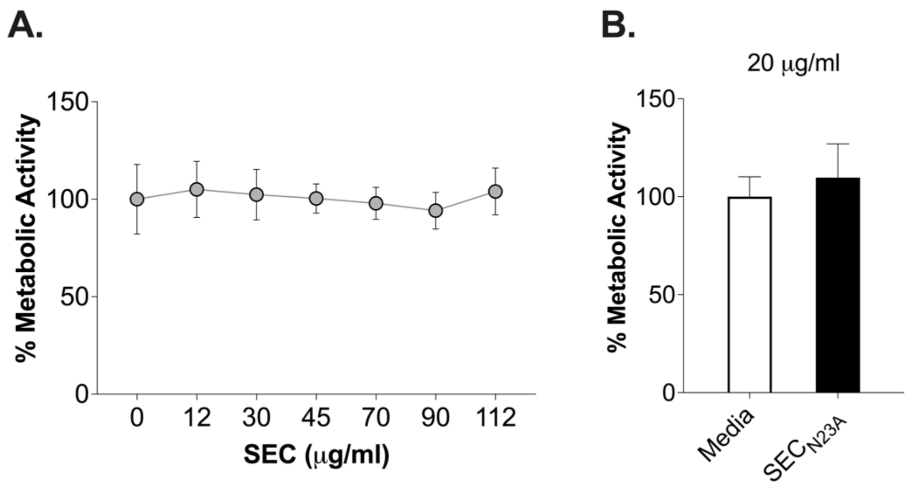


**Figure S6. SEC exhibits no cytotoxicity towards immortalized human aortic endothelial cells (iHAECs).** Percent metabolic activity of iHAECs grown to near confluency on 1% gelatin-coated plates and treated for 24 h with (**A**) increasing concentrations of SEC (12 – 112 μg $mL^{-1}$) or (**B**) $SEC_{N23A}$ at the experimental dose (20 μg $mL^{-1}$). No statistical significance across groups, one-way ANOVA with the Holm-Šídák's multiple comparisons test.

### Supplementary Tables

### Table S1. Strains and plasmids

| Strains | Genotypic Description | Reference |
| --- | --- | --- |
| MW2 | Wild-type *S. aureus* strain from a pneumonia/TSS clinical isolate | 10 |
| MW2Δ*sec* | MW2 with deletion of SEC | 10 |
| MW2Δ*sec*::$P_{sec}$-$SEC_{N23A}$ | Chromosomal complementation of $SEC_{N23A}$ with native MW2 SEC promoter and terminator inserted downstream of MW0058 | This Study |
| MW2Δ*sec*::$P_{sec}$-$SEC_{N23A/F44A/L45A}$ | Chromosomal complementation of $SEC_{N23A/F44A/L45A}$ with native MW2 SEC promoter and terminator inserted downstream of MW0058 | This Study |
| RN4220 | Restriction-deficient cloning strain | 50 |
| **Plasmids** | | |
| pJB38-NWMN2930 | Temp-sensitive shuttle vector for *S. aureus* markerless chromosomal complementation | 46 |
| pUC19 | M13 Cloning Vector | 51 |
| pKK29 | pJB38-NWMN2930 containing $SEC_{N23A}$ with native MW2 SEC promoter and terminator | This Study |
| pKK33 | pUC19 containing $SEC_{N23A}$ with native MW2 SEC promoter and terminator | This Study |
| pKK39 | pUC19 containing $SEC_{N23A/F44A/L45A}$ with native MW2 SEC promoter and terminator | This Study |
| pKK42 | pJB38-NWMN2930 containing $SEC_{N23A/F44A/L45A}$ with native MW2 SEC promoter and terminator | This Study |
| pET25bHSVdelTEV | pET25b vector with HSV tag deleted and a C-terminal TEV cleavage site inserted upstream of the 6xHis tag | This Study |

### Table S2. Mutagenesis Primers

| Target | Oligo | Sequence 5'-3' |
|---|---|---|
| MW2 SEC upstream intergenic region to SEC L27 with N23A mutation in the oligo | pJB38xN23ApromF | GGAGTTTTTTCTGAGTTAAACGATAACCTG AACCTACTGTTGTTAAAAC |
| | promN23AR | AAATATTTCATAGCACCCATCGTACCAGTAA AC |
| MW2 SEC downstream intergenic region to SEC L27 with N23A mutation in the oligo | termN23AF | GGTACGATGGGTGCTATGAAATATTTATATG ATGATCATTATG |
| | pJB38xN23AtermR | TGCAAAATTAAAAATACATCAGGATCAACAC AATTTCATCTAGATGTC |
| $SEC_{N23A}$ with upstream and downstream intergenic regions for insertion into pUC19 | pUC19SECN23AptF | TTGTAAAACGACGGCCAGTGAATTCAACCTG AACCTACTGTTGTTAAAAC |
| | pUC19SECN23AptR | TCTAGAGGATCCCCGGGTACCAACACAATTT CATCTAGATGTC |
| Mutagenesis primers to insert F44A/L45A mutation into $SEC_{N23A}$ | SECF44A/L45Afor | GATAAAGCAGCAGCACATGATTTAATTTATA ACATTAGTGATAAAAAACT |
| | SECF44A/L45Arev | TAAATCATGTGCTGCTGCTTTATCTACAGACT TAACTTTAGTTGCTGATA |
| Screening primers for chromosomal insertion | XNWMN2930F | GCTAATGTGATATATGCAACGCC |
| | XNWMN2930R | GGATGTCCTGTAGCAAATGTTG |
| Sequencing primers for insertion | pJB38XecoRVF | ATGGTATTTGAGCACGTTTAGCGAC |
| | pJB38XecoRVR | CACGGATTGAACACTTTCTACTTAGG |
| Sequencing primer for insertion | CptFseq | GCATAACTTAATTCTTAGAAATAACGTG |

**Table S3. Grading scales and histopathological analyses for the rabbit model of native valve infective endocarditis.**

| | | |
|---|---|---|
| **Heart** | | |
| Histological scoring of myocardial inflammation | | |
| Also noted:<br><br>Vegetation: present (+/-)<br>Myocardial mineralization (+/-) | 0 | No inflammation |
| | 1 | Rare, scattered inflammatory cells within the aortic wall and/or myocardium |
| | 2 | Multifocal bands of inflammatory cells |
| | 3 | Large, coalescing bands of inflammatory cells with necrosis |
| **Kidney** | | |
| Gross pathology scoring | | |
| Also noted:<br><br>Color: red/hemorrhagic (+/-)<br>white/grey/necrotic (+/-)<br>mottled red/grey (+/-) | 0 | No lesions |
| | 1 | Rare, up to 4-5 small (<4mm) multifocal lesions (infarcts) on surface |
| | 2 | Numerous larger (>5mm) multifocal lesions (infarcts) on surface |
| | 3 | Locally extensive to coalescing to diffuse lesions (infarcts) on surface |
| Histological scoring of foci of necropurulent inflammation | | |
| Also scored:<br><br>Ischemic necrosis (infarct) (+/-)<br>Bacterial thrombi (+/-)<br>Intravascular bacterial colonies (+/-)<br>Intralesional bacteria in necrotic foci (+/-) | 0 | No foci |
| | 1 | One foci |
| | 2 | 2 – 5 foci |
| | 3 | >5 foci |
| **Liver** | | |
| Gross pathology scoring | | |
| Also noted:<br><br>Color: white/grey/necrotic (+/-) | 0 | No lesions |
| | 1 | Rare focal streak-shaped lesion/s |
| | 2 | Multifocal to coalescing streak-shaped lesions |
| | 3 | Multifocal streak-shaped and extensive to diffuse |
| Histological scoring of lesions (+/-) | | |
| Centrilobular (CL) vacuolation | | Diffuse to multifocal areas of vacuolated CL hepatocytes |
| Centrilobular (CL) necrosis | | Areas of necrosis in CL hepatocytes |
| Infarcts | | Well demarcated zones of coagulation necrosis, often surrounded by necropurulent inflammation and/or hemorrhage |

**Table S4. RT-qPCR Primers**

| Target Amplicon | Primer Pair | Sequence 5'-3' |
|---|---|---|
| Monocyte chemoattractant protein-1 | MCP-1 For<br>MCP-1 Rev | GCA GAA GTG GGT TCA GGA TT<br>ATT CTT GGG TTG TGG AGT GAG |
| C-reactive protein | CRP For<br>CRP Rev | GTG CTG TCA CCA GAT GAG ATT A<br>TCG CCT TGC ACT TCA TAC TT |
| GM-colony stimulating factor-2 | CSF2 For<br>CSF2 Rev | GTC TCC TGA ACC TGA GTA GAG A<br>GCT CCT GGA GGT CAA ACA TT |
| IL-8 | CXCL8 For<br>CXCL8 Rev | TTT GCC AAG GAG TGC TAA AGA<br>CCA CTC TCA ATC ACT CTC AGT TC |
| Transforming growth factor-β1 | TGF-β For<br>TGF-β Rev | CGT GGA GCT GTA CCA GAA ATA C<br>CAC AAC TCC GGT GAC ATC AA |
| ADAMTS-1 | ADAMTS For<br>ADAMTS Rev | TCA CCA CAG CCC ATG AAT TAG<br>GGA ATC CTG GTT CAC ACC ATT A |
| Angiogenin | ANG For<br>ANG Rev | GGA TAA CTC CAG GTA CAC ACA C<br>CCG TCT CCT CAT GAT GCT TT |
| Serpin E1 | PAI1 For<br>PAI1 Rev | TGG GTG AAG ACA CAC ACA AA<br>CGT TGA AGT AGA GGG CAT TCA |
| Urokinase plasminogen activator | PLAU For<br>PLAU Rev | CCA CAC ACT GCT TCA TTG ATT AC<br>TTC CAC CTC AAA CTT CAT CTC C |
| Acidic FGF | aFGF For<br>aGFG Rev | GGC TCA CAG ACA CCA AAT GA<br>GGC CAA CAA ACC AAT TCT TCT C |
| FGF R-2 | FGFR2 For<br>FGFR2 Rev | CAG CAC TGG AGC CTC ATT AT<br>CAG GTG GTA CGT GTG ATT GA |
| HB-EGF | HBEGF For<br>HBEGF Rev | GGG AAT GGT TTA TGC CCT AGA T<br>CGC TGG GAA CCA TCC ATA TT |
| VEGF-A | VEGFA For<br>VEGFA Rev | ATC AGT TCG AGG AAA GGG AAA G<br>AGG CTC CAG GGC ATT AGA |
| VEGF-C | VEGFC For<br>VEGFC Rev | GCT ACC TCA GCA AGA CGT TAT T<br>ATC GGC AGG AAG TGT GAT TG |
| Amphiregulin | AREG For<br>AREG Rev | ATT GGA CCT CAA TGA CAC CTA C<br>GGA AAT CTC ACT CCC TGA AGA C |
| IGFBP-2 | IGFBP2 For<br>IGFBP2 Rev | CCT CTG GAG CAC CTC TAC T<br>CAG AGA CAT CTT GCA CTG TTT G |
| IGFBP-3 | IGFBP3 For<br>IGFBP3 Rev | CAG AGC ACA GAT ACC CAG AAC<br>GGA CTC AGC ACA TTG AGG AA |
| Pentaxin-3 | PTX3 For<br>PTX3 Rev | CAT TGT TCC TGA GGG AGG AAT C<br>AGT CTC CCA GAG AAG GCT AAT |
| TIMP-1 | TIMP1 For<br>TIMP1 Rev | ATG GACTCT TGC ACA TCA CTA C<br>GGG ATG GAT AAA CAGGGA AAC A |
| TIMP-4 | TIMP4 For<br>TIMP4 Rev | GGT TAC CAG GCT CAG CAT TAT<br>GGC TGA ACG ATG TCA ACA AAC |
| Thrombospondin-1 | TSP1 For<br>TSP1 Rev | CTG GAC TCG CTG TAG GTT ATG<br>GCC AAA GAC AAA TCC AGC ATA G |
| Endoglin | ENG For<br>ENG Rev | GCA GGT GTC AGC AAG TAT GA<br>GAA AGA GAG GCT GTC CAT GTT |
| Endothelin-1 | ET1 For<br>ET1 Rev | GTT GTT CCG TAT GGA CTT GGA<br>GGC TAG CAC ATT GGC ATC TA |
| Angiopoietin-2 | Ang2 For<br>Ang2 Rev | ATC AGG ACA CAC CAC GAA TG<br>CAT CCT CAC GTC GCT GAA TAA |
| Beta-actin | ACTB For<br>ACTB Rev | GGA TCA GCA AGC AGG AGT ATG<br>AGA AAG GGT GTA ACG CAA CTA A |
